# Foundation Models Improve Perturbation Response Prediction

**DOI:** 10.64898/2026.02.18.706454

**Authors:** Elijah Cole, Geert-Jan Huizing, Sohan Addagudi, Nicholas Ho, Euxhen Hasanaj, Merel Kuijs, Toby Johnstone, Maria Carilli, Alec Davi, Caleb Ellington, Christoph Feinauer, Pan Li, Romain Menegaux, Shahin Mohammadi, Yanjun Shao, Josiah Zhang, Emma Lundberg, Le Song, Ziv Bar-Joseph, Eric P. Xing

## Abstract

Predicting cellular responses to genetic or chemical perturbations is a long-standing goal in biology. Recent applications of foundation models to this task have yielded contradictory results regarding their superiority over simple baselines. We conducted an extensive analysis of the utility of pretrained embeddings (including but not limited to biological foundation models) for perturbation response prediction, covering over 600 embedding-task pairs. While some foundation models fail to outperform simple baselines, others yield consistent improvements. However, the strongest pretrained embeddings are those derived from prior knowledge. We also developed methods for integrating multiple pretrained embeddings to make better predictions. Our results confirm that pretrained embeddings can enable more accurate simulations of cellular behavior.

**Code and Data:** https://github.com/genbio-ai/foundation-models-perturbation

## 1 Introduction

Predicting the outcome of cellular perturbations is a long standing challenge in molecular biology [1–3]. Typically these perturbations are either genetic modifications (e.g. knocking out or over-expressing one or more genes [4]) or chemical treatments (e.g. exposing cells to a small molecule [5]). Readout types for perturbation experiments include cell death [6, 7], cell growth [8], and imaging [9]. One of the most interesting readouts is gene expression [10], which measures the impact of a perturbation on some or all genes in a cell. Initial perturbation response prediction models focused on bulk expression [11] while more recent efforts consider single cell perturbation responses [12].

There are several reasons why perturbation prediction is an important question in biology. First, the ability of models to predict the impact of a perturbation in a specific context (cell type, tissue, disease state, etc.) suggests that such models may have learned something useful about the underlying molecular networks that are active and repressed in that context [13]. A model that accurately predicts perturbation responses in diverse settings may have learned to account for both the activity of the perturbed gene (or the target of the small / large molecule) and the interactions between that gene / protein and other proteins and pathways in the cell. Such knowledge is one of the main goals of systems biology. In addition to demonstrating that a useful representation of cell state and interactions has been learned, such analysis is also a key part of most drug discovery and design processes. The ability to accurately predict the impact of a specific treatment at the cellular level can save time and effort when identifying new interventions for diseases [14]. In addition, such models can provide information not just on the direct effects of a treatment but also alert researchers to potential indirect or side effects, which in many cases are only discovered very late in the drug discovery process [15]. Accurate perturbation response prediction would be impactful for both basic and applied biology research, which has motivated ongoing interest in developing such models.

Early work on perturbation modeling aimed to predict drug mechanism of action [16], estimate sensitivity to drugs [17], or model gene regulatory networks with differential equations [1, 18, 19]. As the available data grew, perturbation response prediction problems were studied using statistical learning methods including Elastic-Net [6, 20, 21], neural networks [14], and matrix factorization models [22, 23]. Early applications of deep learning include Dr.VAE [24] and scGen [25]. These initial deep learning frameworks have been extended in several directions. CPA [26] enabled the prediction of combinatorial perturbation effects. GEARS [27] incorporated knowledge graphs and allowed for the prediction of perturbations not seen during training. Most recently, transformer-based architectures have been adapted from the mainstream ML community, leading to models like Geneformer [28], scFoundation [29], and scGPT [30], which pretrain on massive single-cell atlases and provide schemes to use the FM for perturbation response prediction. These recent papers have claimed to achieve strong results using such methods for perturbation analysis. However, other recent publications [31, 32] take a contradictory view, arguing that FMs do not improve on simple baseline methods. Concurrent work has argued that new, carefully defined metrics are required to separate complex methods from naive baselines [33].

Here we perform a comprehensive study to resolve this issue by evaluating the use of pretrained embeddings, including but not limited to those from biological FMs, for perturbation prediction. We evaluate many embeddings from FMs and compared them to simple baselines, classical ML methods, and recently state-of-the-art methods across several prediction tasks for different perturbation types. Inspired by practices in mainstream machine learning, we take an embedding-centric view. Unlike many prior works which consider only scRNA-seq FMs, we generate perturbation embeddings using FMs trained for modalities like DNA sequences [34], protein sequences [35], protein structures [36], and small molecules [37]. Our analysis indicates that while some FMs indeed perform no better than simple baselines, others consistently outperform them. However, pretrained embeddings derived from prior knowledge tend to perform best. We further show that integrating collections of pretrained embeddings using attention-based fusion can lead to further improvements.

## 2 Results

### 2.1 Embeddings vary dramatically in utility for genetic perturbation prediction

Prior studies have claimed that there is no clear benefit from using FMs for predicting the effect of unseen perturbations, arguing that simple baselines (e.g. linear models with embeddings derived from PCA) perform as well or better (e.g. [31, 32, 38]). We would draw similar conclusions if we considered only the dataset from Norman et al. [4], on which we find that no embedding improves meaningfully on Train Mean (Figure A1).

However, if we consider the much larger Essential dataset [39, 40] (four CRISPRi screens, *>*2000 perturbations each) we readily observe that many embeddings, including some from biological FMs, do outperform simple baselines. We first note that, unlike in Norman, there are clear differences between embeddings that are consistent across cell lines (Figure 1(a, b)). While some embeddings perform no better than simple baselines like Train Mean and PCA, most embeddings we study beat these simple baselines to varying degrees. These successful embeddings include some from scRNA-seq FMs, protein sequence FMs, and many embeddings derived from prior knowledge. While the FM embeddings are not the top performers, they do outperform the naive baselines. To fortify this finding, we re-evaluate our embeddings in the formulation of Ahlmann-Eltze et al. [32] using their code, splits, and metrics. Figure A9 shows that our embedding rankings are similar in their setting. Notably, many of our embeddings outperform all models evaluated in Ahlmann-Eltze et al. [32].

**Fig. 1.**
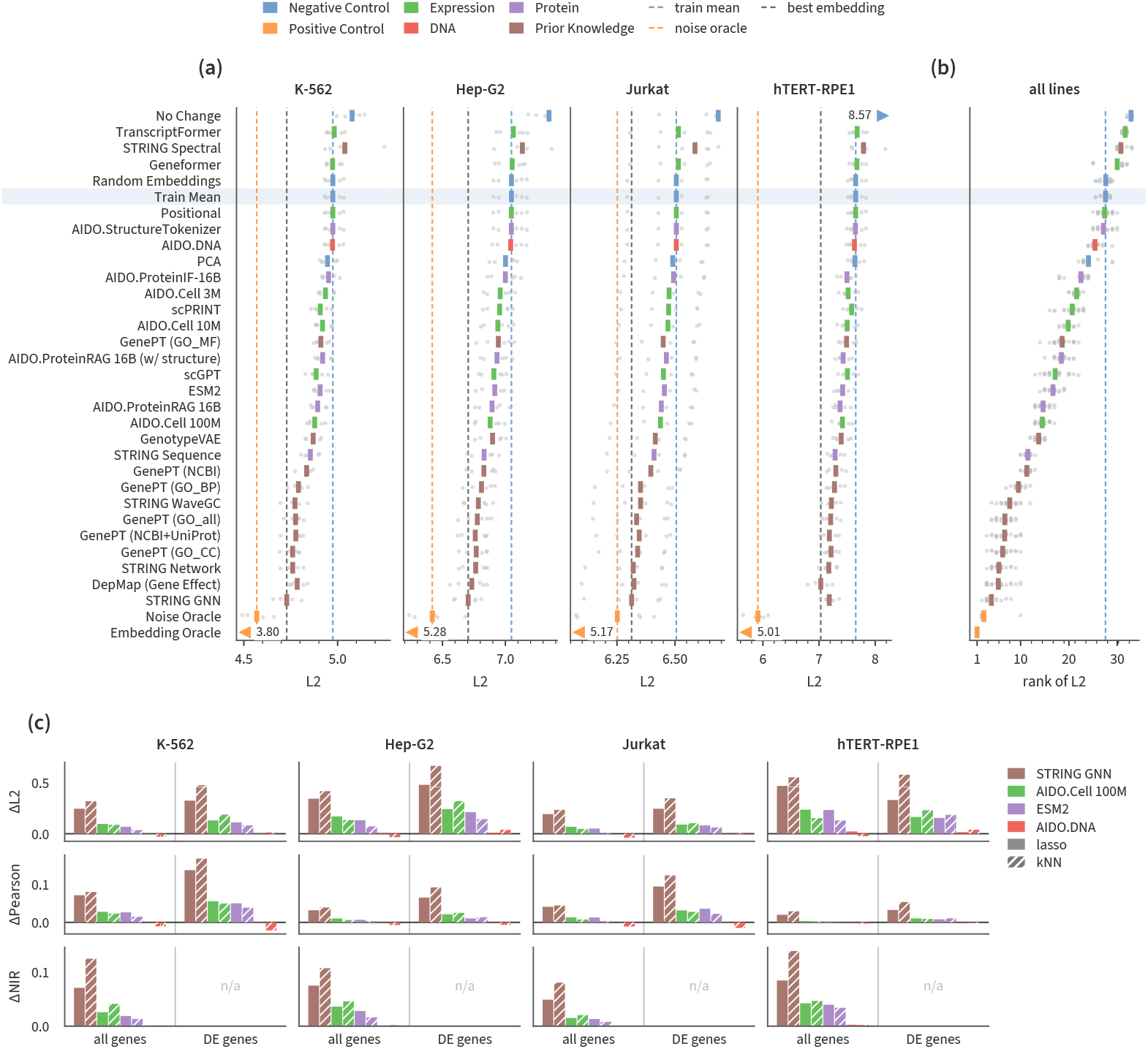
Many embeddings, led by the prior knowledge family, consistently improve on the train mean in the four Essential screens. Out-of-fold LFC prediction with lasso on frozen gene embeddings in the four Essential screens. (a) L2 per method and cell line (lower is better). Gray dots are folds, ticks are the mean over folds, and triangles represent off-axis means. Dashed lines mark the train mean, the noise oracle, and the best embedding. (b) Rank of L2 in each cell line and fold. Rows are sorted by mean rank, best at the bottom. (c) ΔL2, ΔPearson, and ΔNIR over the train mean for one embedding per family, with lasso (solid) and kNN (hatched). NIR has no DE-gene version (Supplementary Section B.1).

Interestingly, we find that when we rank embeddings by performance (Figure 1(b)), they tend to be roughly ordered by data modality. The strongest embeddings are typically those derived from prior knowledge like interactome data (e.g. STRING GNN), gene effect screens (e.g. DepMap), or textual descriptions of genes (e.g. GenePT). The best-ranked models after the prior knowledge group is biological foundation models trained on expression or protein data. Among these, the best are STRING sequence, AIDO.Cell 100M, AIDO.ProteinRAG 16B, ESM2, and scGPT. Some foundation models, such as Geneformer and TranscriptFormer, rank below the train mean. These broad rankings are consistent with findings in Littman et al. [41]. This pattern does not seem to be a property of the specific supervised task we are considering, since we see similar patterns in a discrete DEG classification formulation (Figure A3). Further, a direct analysis of the pairwise similarity between embedding sources shows similar modality-driven clustering patterns (Figure 2). In Figure 1(c), we use four family representatives (STRING GNN, AIDO.Cell 100M, ESM2, AIDO.DNA) to show that this general pattern is robust to metric, prediction head type, and evaluation gene set. We further show that this pattern extends to other genetic perturbation screens (Figure A10) and more challenging splits based on gene family holdouts (Figure A12). Embedding rankings are highly stable across different metrics and evaluation gene sets when perturbations carry replicable signal (Figure A13). Finally, we use GO gene sets to analyze the types of biology each embedding predicts best (Section 2.5).

**Fig. 2.**
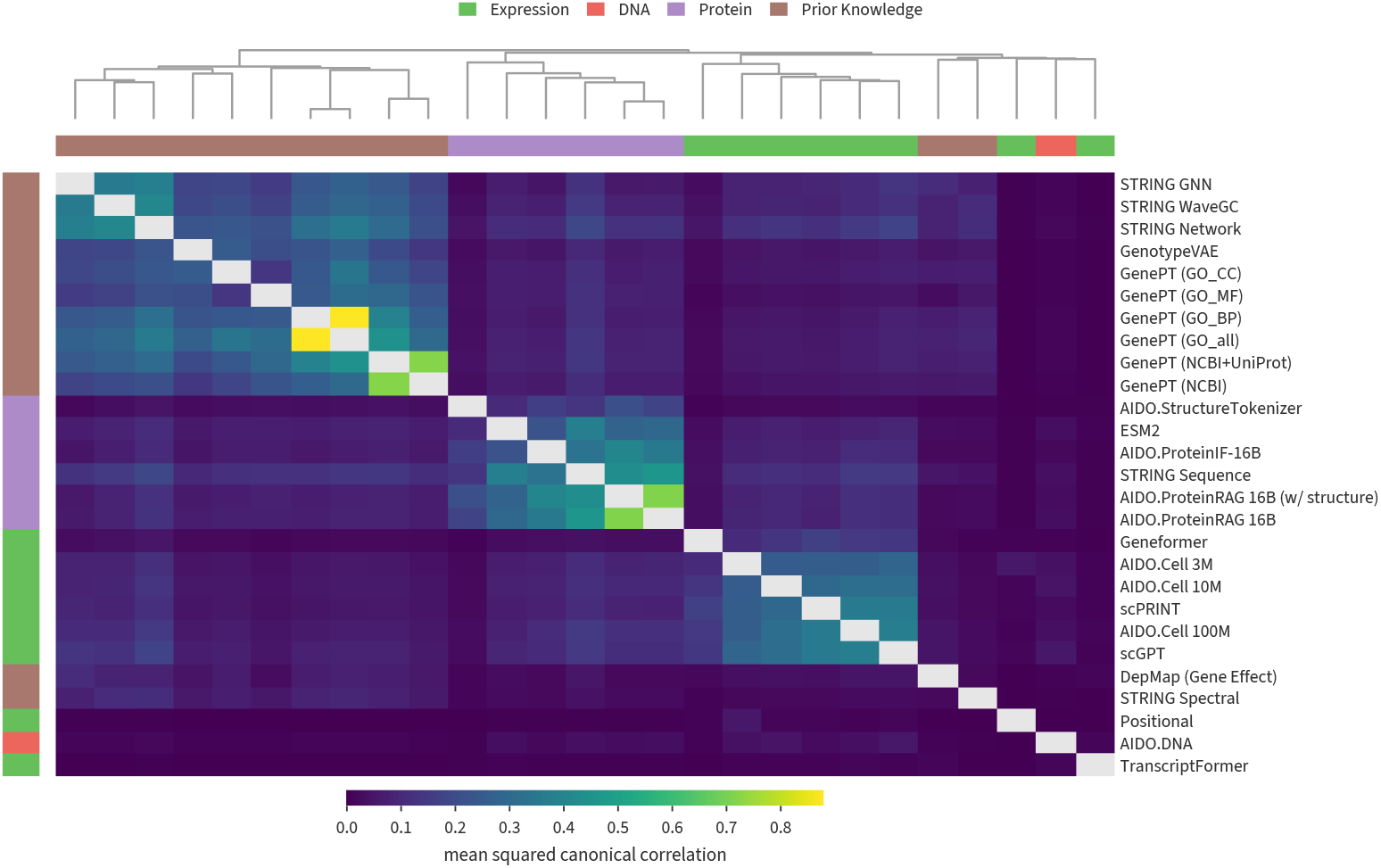
Gene embeddings cluster by family. Pairwise similarity between gene embeddings on their shared genes, as the mean squared canonical correlation of their top principal components (Supplementary Methods). Cell-line-specific embeddings are the K-562 versions. Rows and columns are ordered by hierarchical clustering. Color strips mark the family, and the diagonal is omitted.

The benefit of pretrained embeddings depends on how reproducible a perturbation’s response is. Across screens, gains over the train mean appear only for perturbations with replicate SNR above about 0.5 (Figure A10). Such perturbations make up 76–94% of the Essential screens but 39% (HEK293T), 17% (HCT116), and 3% (iPSC) of the others, which explains why aggregate gains there are small. Benchmarks that differ in this composition can therefore reach opposite conclusions about the same models if interpreted only at the aggregate level.

Some of our embedding-based predictors are surprisingly strong. On the K-562 cell line the best embedding closes most of the gap between Train Mean and Noise Oracle (a positive control that has access to the test labels). However, different types of positive control vary substantially in terms of how much remaining room for improvement they suggest (Figure A14).

### 2.2 Complex embedding translation methods do not significantly outperform simple ones

The benchmarking results presented in Figure 1 use Lasso or kNN for translating embeddings to average expression change vectors (Methods). However, there have been several suggestions in the literature for a different formulation based on modeling populations of single cells [42–44].

To test these ideas, we used our Essential benchmark to evaluate several distribution prediction methods which predict distributions of perturbed cells given a perturbation embedding and a collection of control cells. Instead of evaluating published perturbation prediction systems, we study simple implementations of the underlying generative models: Latent Diffusion [45], Flow Matching[46], and Schrödinger Bridge [47]. This allows us to decouple the embeddings used for perturbation conditioning from the generative model itself, enabling clear comparisons to the other methods we consider in this work.

We evaluate these methods in the LFC regression formulation by reducing their predicted distributions to their means. Full details can be found in Supplemental Methods. In addition, we evaluated a published GNN-based perturbation model (GEARS). As can be seen in Figure A6, none of these advanced methods outperform kNN combined with our strongest embedding. Importantly, Latent Diffusion, Flow Matching, and Schrödinger Bridge start from the exact same embedding as kNN. These methods are significantly more computationally expensive, so they received less extensive hyperparameter tuning than kNN (Methods) despite consuming far more GPU-hours. Under a generous but fixed budget, these advanced methods did not significantly improve over kNN in our setting.

Our main focus is on directly using embeddings from frozen FMs (Methods). While FMs can learn powerful representations of biology through large-scale pretraining, fine-tuning may further improve their performance [28, 30, 48]. A full exploration of fine-tuning is beyond the scope of this work, but we conduct a preliminary exploration by fine-tuning two of the best performing models from different categories (AIDO.Cell (3M) and STRING GNN) on the LFC regression task on Essential (Figure A7). Our findings were mixed, and we leave further exploration of fine-tuning to future work.

This work is primarily concerned with predicting the effects of unseen perturbations in seen cell lines. Recent works like STATE [49] consider a different context: predicting seen perturbations in cell lines for which limited data (few shot) or no data (zero shot) is available. We thus also compared an embedding-based MLP approach against STATE on the Essential (Figure A8) and Tahoe (Figure A8) datasets. The embedding-based MLP performs similar to or better than STATE in 4 out of 6 dataset-metric combinations we evaluate, improving on STATE by up to 64%. However, it does not seem that the embeddings themselves are the primary reason for this performance gain. In an ablation where we replace the embedding with a one-hot encoding of the perturbation, the performance of our MLP does not change much in either dataset for most metrics.

### 2.3 Pretrained embeddings can also improve small molecule perturbations predictions

For genetic perturbation datasets like Essential, perturbations can be naturally represented by embeddings of genes or proteins. The picture is more complex for chemical perturbation datasets (Figure B16). The traditional way to embed a small molecule is to use the SMILES or graph representation of the molecule to produce an embedding, either with a traditional feature encoding (e.g. Morgan fingerprints [50, 51]) or a transformer-based encoder (e.g. ChemBERTa [37]). An alternative is to represent a molecule by embedding its target (either the gene or a protein product) with a pretrained model. However, this requires additional information that is not always available. A third approach is to take textual descriptions of the molecule from a database (e.g. ChEMBL) and use an LLM to reduce that text to a single embedding. Again, the utility of this depends on what information is available in databases.

We compared several traditional and FM based methods using the Tahoe-100M dataset (Figure A4). These were evaluated in a benchmarking setting similar to our Essential results. As before, the task is to predict the effect of unseen perturbations. In the LFC regression formulation, we find that no embedding convincingly outperforms the negative controls. However, the DEG formulation of Tahoe shows that some embeddings clearly outperform negative controls (Figure A4). As a category, target-based embeddings performed best - simple target encodings, AIDO.Cell 100M, and scPRINT all performed well. The best-performing molecular structure fingerprint (ECFP:2) fingerprint was not far behind. The targets are LLM suggestions whose accuracy we do not assess (Methods). We simply test their utility in our downstream evaluations. Note that better-studied molecules likely receive better suggestions. The SMILES-based foundation models (with the exception of MiniMol) tended to under-perform, possibly due to the fact that they have mostly been developed with chemical (not biological) applications in mind. We provide complementary results for Sciplex-3 in Figure A5.

### 2.4 Integrating diverse embeddings can further improve performance

All of the results presented so far have used a single perturbation embedding at a time to predict perturbation effects. However, objects like small molecules, genes, and proteins are complex and multifaceted. Any given representation may capture some aspects and neglect others. It therefore makes sense to integrate information from different models and modalities. In this section, we introduce and evaluate an attention-based model for integrating embeddings and predicting perturbation response (Methods).

Our results for Essential in Figure 3 show that multimodal fusion always improves over the best unimodal embedding for Essential, and that the more complex Fusion (Full) model improves over the Fusion (Simple) model. For two cell lines (Jurkat, K-562), the fusion result matches or exceeds the Noise Oracle positive control. Fusion results for the other two cell lines are also better than the single strongest embedding. Figure A11 replicates these findings for a Perturb-Seq screen in iPSCs, for the perturbations with replicable signal.

**Fig. 3.**
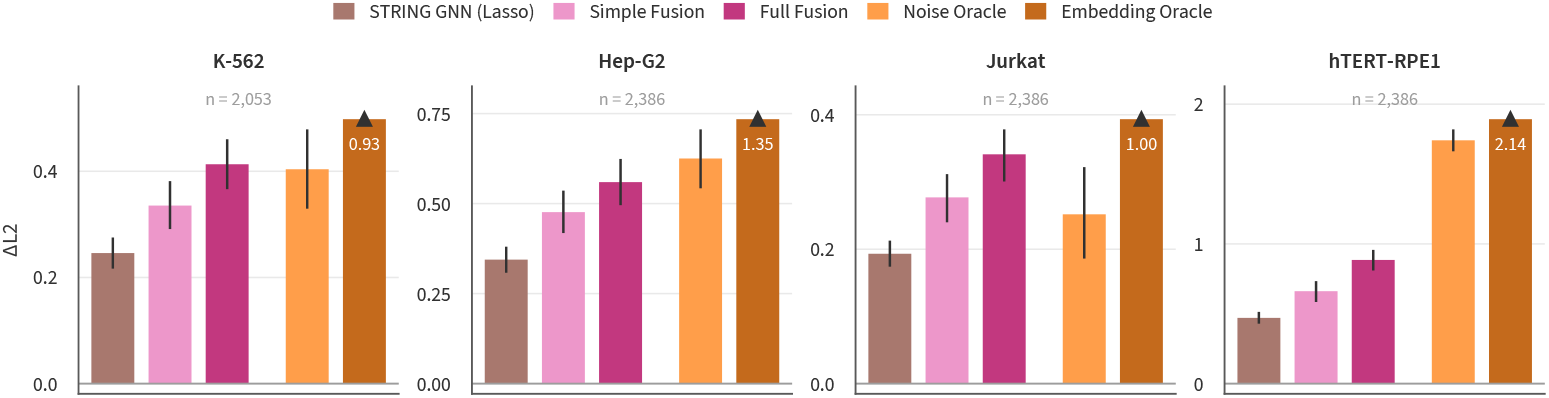
Fusing many embeddings improves on the best single embedding in every Essential screen, and it reaches the noise oracle on K-562 and Jurkat. ΔL2 over the train mean on the four Essential screens for lasso on STRING GNN, the simple and full fusion models, and the noise and embedding oracles. Both fusions combine frozen gene embeddings (all pretrained embeddings from Figure 1) and are trained jointly on the four lines. Each panel uses the perturbations that every bar scores, with the number of perturbations in gray. Error bars are 95% bootstrap CIs over perturbations (Supplementary Section B.1). The embedding oracle runs past the axis, and its value is printed on the bar.

However, the fusion approach is not universally helpful. We do not find any benefit for the DEG prediction formulation of Essential (Figure A3), Sciplex (Figure A5), or Tahoe (Figure A4).

### 2.5 Analysis of gene set predictions identifies processes that are easy and hard to predict

In addition to our general conclusion about the advantage of FMs that are based on a curated set of low throughput experiments, we have also studied the impact of the biological processes on the accuracy of the predictions.

We first analyze the extent to which process predictability depends on the embedding, the dataset, or the interaction between them (Figure A15). There are two ways we can assign scores to gene sets. First, we can score each gene set in terms of how well their corresponding perturbations are predicted (“perturbation lens”). Second, we can score each gene set in terms of how well those genes are predicted as read-outs (“readout lens”). For each case, we perform a two-way ANOVA to decompose the observed variance into an embedding-driven component and a dataset-driven component. As Figure A15(a) shows, the perturbation lens depends more on the type of embedding-used, as we have seen for the general prediction scores. On the other hand, Figure A15(b) shows that the readout lens depends more on the dataset. In other words, embeddings tend to agree on which readout genes are predictable, but which genes those are varies by dataset. Mitotic spindle regulation and translational elongation vary most by cell line. Figure A15(c) further breaks down the variance within each cell line, with components for the biological process, the embedding, and the residual (“specialization”).

In Figure 4 we select four representative embeddings and perform gene set enrichment analysis to identify the gene sets that each embedding predicts best. Ribosome biogenesis is significant for STRING-GNN in all six cell lines, and for AIDO.Cell and ESM2 in five cell lines. Mitochondrial ATP synthesis and metaphase chromosome alignment are only significant for AIDO.Cell and ESM2, while mitochondrial translation is significant only for STRING-GNN. AIDO.DNA has no enriched processes, likely because the prediction quality is generally poor (Figure A10).

**Fig. 4.**
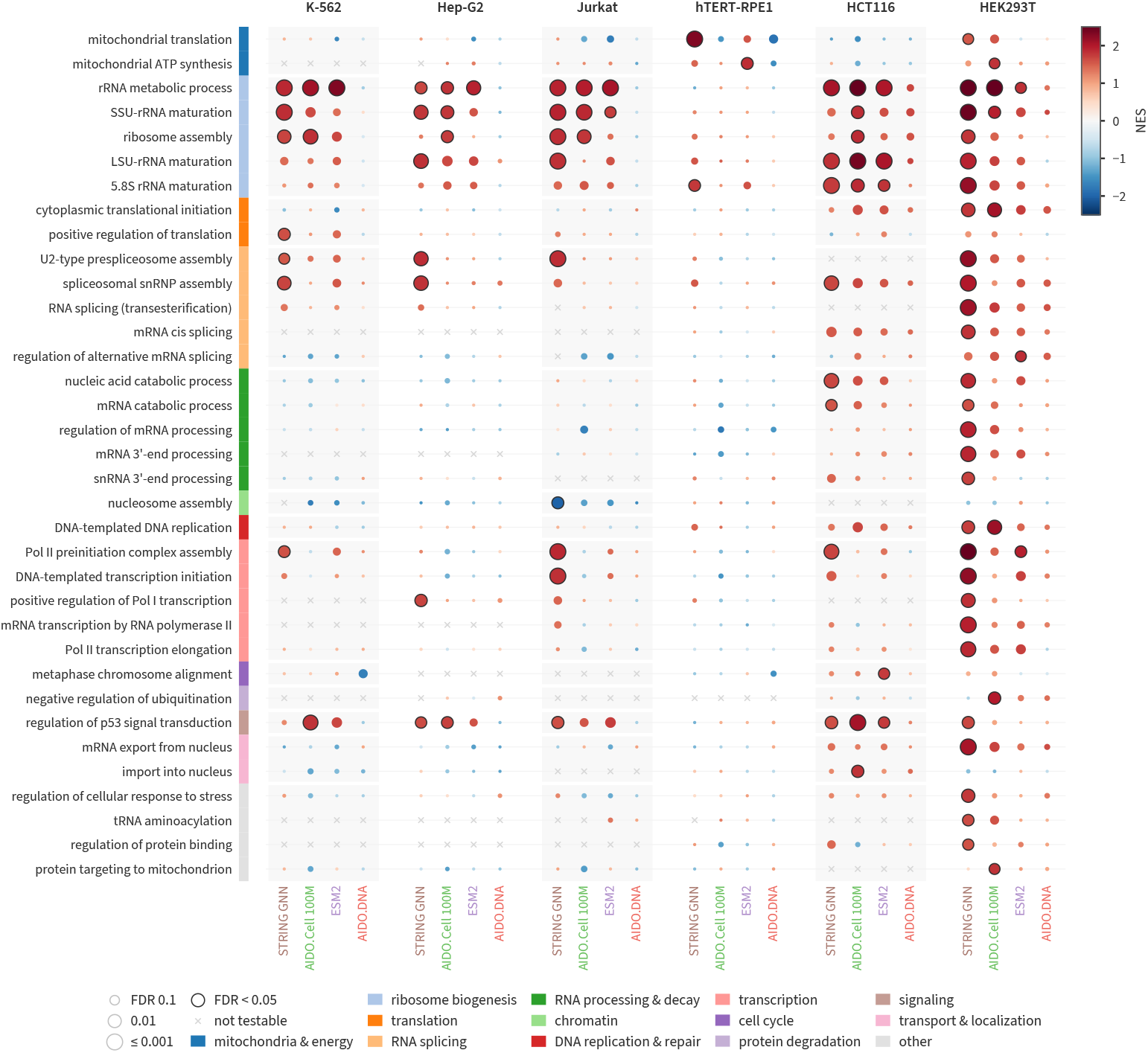
Gene sets predicted best by each of four representative embeddings and 6 cell lines, as determined by preranked gene set enrichment analysis. Dots are colored by the normalized enrichment score (NES) and sized by false discovery rate (FDR). For ranking, perturbations are scored using the out-of-fold *R*^2^ relative to Train Mean (Section B.1). We start with 144 gene sets and show every gene set that is significantly enriched for at least one (embedding, cell line) combination.

## 3 Discussion

The use of FMs for perturbation prediction has received a lot of attention over the last few years with contradictory findings. Here we performed a comprehensive analysis of embedding-based perturbation response prediction for both genetic and small molecule interventions. As we showed, while some FMs fail to improve over simpler methods for this task, others provide significant gains. Embeddings based on interaction networks and other prior knowledge sources performed particularly well. In addition, we show that some biological processes are easier to predict than others for all embedding methods.

We also observe that predicting genetic perturbations is an easier task than predicting the impact of a small molecule intervention. This is true even when the target is (assumed to be) known for the small molecule. This is may be due to the higher specificity of genetic perturbations (i.e. small molecules are more likely to target multiple proteins or pathways). In addition, the search space of possible small molecules is much larger (estimated to be ∼10^60^ [52]) than the one we use for genes (which is limited to ∼2 ×10^4^). Further, we usually have much less information on the underlying interaction networks for small molecules which, as we showed, provide the most valuable information for perturbation prediction.

While we observed that specific embedding families outperformed other families, integrating several different types of embeddings often improved predictions for genetic perturbation prediction. This is consistent with prior works that have experimented with combining two embeddings at a time [53, 54]. We have tested various strategies for integrating many embeddings and shown that in some cases, embedding fusion is sufficient to match positive controls that have access to the test labels. In the small molecule setting, fusion of FMs did not improve DEG prediction performance on either Sciplex or Tahoe, but Simple Fusion improved LFC regression on Sciplex. We speculate that the weaker fusion results are due to the weak performance of most individual embeddings and the potential to overfit when learning fusion models with a smaller number perturbation-context pairs.

Our study provides strong support for the use of pretrained embeddings for perturbation prediction. Still, there are many open questions and potential research directions to improve and follow up on the results we presented. One such direction relates to the type of data that is most useful for perturbation prediction. We found that, for genetic perturbations, interactome data seems to be the most valuable. This should be taken into account by organizations that are aiming to study and develop virtual cell models. Specifically, this result indicates that data related to interactions in different contexts (cell types, stages of development, diseases, etc.) would be a valuable addition to public data sources, perhaps more than single cell expression data. For small molecule perturbations, we have consistently found that off-the-shelf small molecule embeddings perform very poorly on our small molecule benchmarks. Many of these models were trained to predict chemical properties, which may not connect closely to biological function. The field seems to lack a standard foundation model for small molecules that reflects biological function. Such a model may provide large benefits for predicting small molecule perturbations.

Another way to improve the results is fine-tuning. We have briefly explored fine-tuning individual models with mixed results (Figure A7), but a more thorough analysis should be conducted in the future. Fine-tuning of integrated multimodal models may improve performance further, but this is a much more challenging and computationally demanding task. It remains to be seen if jointly fine-tuning multiple pretrained models in a fusion framework can lead to further improvements. The risk of overfitting would be high without additional training data. To conclude, our comprehensive analysis indicates that pretrained embeddings are pushing the boundaries of our ability to accurately predict perturbations. Larger datasets and better FM methodologies may lead to increasingly accurate models for this task.

## 4 Methods

### 4.1 Formulation of Log Fold-Change Regression

#### 4.1.1 Notation

We start with a collection of *N* scRNA-seq profiles from a perturbation experiment in which K different perturbations *P*_1_, …, *P*_*K*_ are tested. If we measure *G* genes for each cell, our observed expression matrix can then be written *X* ∈ ℝ^*N×G*^. Let *X*_*i*_ denote the *i*th row of *X*. We assume each row *X*_*i*_ has been total count normalized, scaled by 10^4^, and log1p transformed. Let *S*^ctrl^ be the set of indices for rows of *X* corresponding to control cells and let 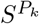 be the set of indices for rows of *X* corresponding to cells receiving perturbation *P*_*k*_. Thus, 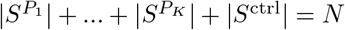. Finally, we assume our *N* cells are partitioned into *M* batches based on some experimental condition. To describe this, let *S*(*b*) be the set of indices of cells belonging to batch *b*. This means that 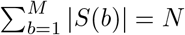. Let

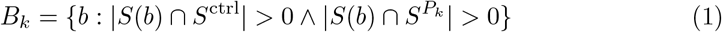

denote the set of batches containing both control cells and cells that received perturbation *P*_*k*_.

#### 4.1.2 Definition of Ground Truth Labels

Our focus in this work is predicting the average treatment effect (ATE) of unseen perturbations. To account for batch effects, we weight the contribution of each batch equally. The result is a “batch aware” average treatment effect (BA-ATE). We use two variants, depending on whether the dataset’s batches are typically large (≥100) or small (*<* 100).

First, define the average control expression in batch *b* as

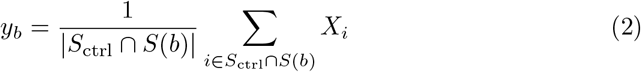

where *y*_*b*_ ∈ℝ^*G*^. Similarly, define the average expression under perturbation *P*_*k*_ in batch *b* as

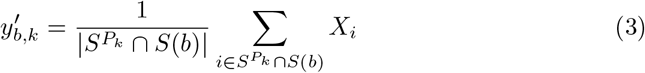

where 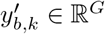.

Throughout the paper we refer to the BA-ATE as “log fold-change” (LFC). This is an abuse of terminology, but a useful one since BA-ATE is very similar in spirit. The differences are that BA-ATE (i) includes a batch adjustment, (ii) uses a log1p transform instead of a log transform, and (iii) performs averaging after the log1p transform.

##### Datasets with large batches (Tahoe, Sciplex, Norman)

Large batches allow us to accurately determine perturbation effects using perturbed and control cells from the same batch. We define the BA-ATE for large batches as

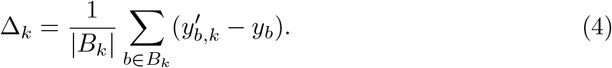

##### Datasets with small batches (Essential)

When batches are small, estimates of per-batch control expression are likely to be noisy. In this case, we first use all batches (whether or not there are matched perturbed cells) to estimate the global control expression for gene g:

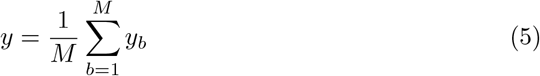

where *y* ∈ ℝ^*G*^. Then we compute the BA-ATE for small batches as

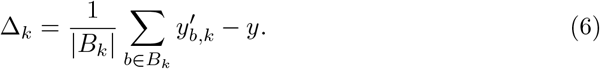

##### iPSC

With about 7 cells per perturbation and lane, batches are small. Δ_*k*_ is the mean expression over all cells of *P*_*k*_ minus a control mean. Each sublibrary has its own control mean, over its non-targeting cells. The control for *P*_*k*_ averages these means, weighted by the number of *P*_*k*_ cells in each sublibrary.

##### Xaira

Batches hold about one cell per perturbation, so we pool control cells over batches. Δ_*k*_ is the mean expression over all cells of *P*_*k*_ minus the mean over all non-targeting cells.

#### 4.1.3 Metrics

Let Δ_*k*_ ∈ ℝ^*G*^ be the vector of per-gene treatment effects for perturbation *P*_*k*_. Let 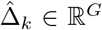 be some model prediction for the same quantity. Our default performance metric is the simple L2 error, averaged over perturbations:

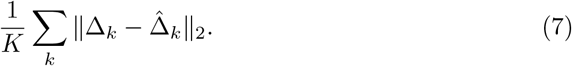

All metrics, aggregation methods, and confidence interval calculations are detailed in Section B.1.

### 4.2 Formulation of Differentially Expressed Gene (DEG) Classification

#### 4.2.1 Notation

We begin with the same perturbation experiment setting and notation described in the LFC regression problem formulation in Section 4.1.1. We use parallel notation where possible (e.g. Δ_*k*_ for the effect of perturbation *P*_*k*_) since the meaning is clear from context.

#### 4.2.2 Definition of Ground Truth Labels

In this problem formulation, we aim to predict the differential expression status of each gene instead of the exact change in expression. We account for batch effects and weight the contributions of each batch equally.

For batch *b*, the differential expression results for perturbation *P*_*k*_ are computed as

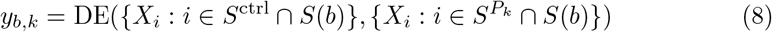

where the DE operator performs an independent differential expression test for each gene using the control cells {*X*_*i*_ : *i* ∈ *S*^ctrl^ ∩*S*(*b*)} and the perturbed cell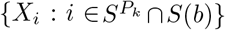. This results in a vector *y*_*b,k*_∈{− 1, 0, +1}^*G*^, indicating whether each gene is downregulated (−1), upregulated (+1), or not significantly different (0).

##### Datasets with large batches (Tahoe, Norman, Sciplex)

After computing per-batch differential expression vectors *y*_*b,k*_, we produce a final ground truth perturbation effect Δ_*k*_ ∈ {−1, 0, +1} ^*G*^ for perturbation *P*_*k*_ by performing majority vote over batches.

##### Datasets with small batches (Essential)

For Essential, the batches are too small to obtain meaningful per-batch differential expression results. We therefore ignore batch structure when performing the differential expression testing, comparing all perturbed cells to all control cells:

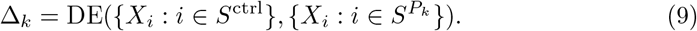

##### Implementation details

Differentially expressed genes were called using a Welch t-test with Benjamini-Hochberg multiple testing correction (FDR *<* 0.05) across genes for each perturbation. The perturbed group’s variance is floored using the control group’s variance. We discard perturbations with ≤30 cells. For iPSC, we use sublibrary-matched controls and require an absolute mean difference in log1p expression greater than 0.1. For all other genetic perturbation datasets, the control population is pooled over batches. For Sciplex and Tahoe, we run the test for each (perturbation, plate) combination with *>* 20 cells and make final DEG calls by majority vote.

#### 4.2.3 Metrics

Our metric is based on the F1 score and defined as follows. We first compute an F1 score for each gene and class (up, down, no change). We obtain an F1 score per gene by averaging over the classes present in each gene’s label set. Then we average these gene-level F1 scores over all genes that are DE at least once in the held out perturbations for that fold. All metrics, aggregation methods, and confidence interval calculations are detailed in Section B.1.

### 4.3 Data

Here we describe the perturbation datasets used for embedding benchmarking.

All datasets are normalized as in Section 4.1.1 (total counts to 10^4^, then log1p) before all downstream calculations. Note that the Essential and Tahoe datasets undergo special preprocessing for the cross-context comparison with STATE (Supplemental Methods).

#### 4.3.1 Norman

The Norman dataset is a genetic perturbation dataset (CRISPR activation, i.e. overexpression of the target gene) in one cell line, K562 [4]. The genes selected for perturbation were known to be involved in the regulation of cell growth. The dataset contains both single and dual perturbations. We use only the 105 single gene perturbations. This subset of the Norman dataset has 8 large batches (13k+ cells each), which are defined by GEM group.

#### 4.3.2 Essential

The Essential dataset is a genetic perturbation dataset (CRISPR knockdown) of over 2000 “essential” genes per cell line for four cell lines (2387 for Hep-G2, Jurkat, and hTERT-RPE-1; 2054 for K-562.). There are 964451 single cells profiled. It has been compiled from two independent publications from the same lab [39, 40]. The Essential dataset has 56 small batches, which are defined by GEM group. There are 133784 unique (batch, perturbation) pairs, so on average there are 7.2 cells per perturbation and batch.

#### 4.3.3 iPSC

The iPSC dataset is a genome-scale CRISPRi screen in the KOLF2.1J induced pluripotent stem cell line [55]. Most genes are targeted by three guides. We use 11,686 perturbed genes and 2,733,495 cells, of which 151,401 carry non-targeting guides. The median perturbation has 224 cells. Cells were profiled in 90 lanes, grouped into three sublibraries that each have their own non-targeting cells. The readout is 17,701 protein-coding genes. We use iPSC for the SNR-stratified analyses, the gene-family split, and metric agreement.

#### 4.3.4 Xaira

The Xaira datasets are genome-scale CRISPRi screens with dual guides in HCT116 and HEK293T cells [56]. We use 18,293 (HCT116) and 18,311 (HEK293T) perturbed genes, with a median of 150 and 200 cells per perturbation. Cells were profiled in 109 (HCT116) and 223 (HEK293T) GEM samples, about one cell per perturbation and sample. The benchmark readout is 19,401 protein-coding genes. Replicate SNR (Supplementary Section B.9) is computed on all 38,606 measured genes. We use Xaira for the SNR-stratified analyses and the gene set analyses.

#### 4.3.5 Sciplex

SciPlex3 [5] is a single cell chemical perturbation dataset. The raw expression matrix contains 799,317 single cells and 110,983 gene features, reflecting the full unfiltered set of detected transcripts. SciPlex3 profiles three human cancer cell lines: A549 lung adenocarcinoma, MCF7 breast adenocarcinoma, and K562 chronic myelogenous leukemia. Cells are treated with 188 distinct small molecule drugs as well as controls. Each drug is applied at four concentration levels: 10 nM, 100 nM, 1000 nM, and 10000 nM. In this work, we restrict the analysis to protein coding genes only. After filtering, the dataset contains 18,590 protein coding genes, which are used for all downstream modeling and evaluation. We further restrict all experiments to the highest dose level of 10000 nM for each drug. This dataset is organized into 52 large sized batches, consisting of 154.4 cells on average for each (plate, drug, dose) combination (of which there are 4940).

#### 4.3.6 Tahoe

The Tahoe-100M dataset is a chemical perturbation dataset generated by Tahoe Therapeutics [57]. It contains ∼100M cells and measures ∼60k genes. The cells are from 50 cell lines and profile 379 drugs, each at 3 dose levels. We used the highest dose - 5 uM - in our experiments. We excluded 5 cell lines (CVCL 1571, CVCL 1531, CVCL 1715, CVCL 1577, CVCL 1716) because their control cells had unusually low total counts. We also removed any compounds without SMILES string representations. The Tahoe dataset has 14 large batches, which are defined by the plate on which the cells were cultured. There are 15932 unique (plate, drug, dose) triplets, with an average of 6317.4 cell each.

### 4.4 Compute

Many of the methods used, studied, and tuned in this work required significant compute resources. Most notably, training and evaluating some of the advanced methods for single cell expression matching (Latent Diffusion, Schrödinger Bridge) required ∼1000 GPU-hours each. Other parts of the study that relied on extensive compute include the fine-tuning of expression and interactome FMs and learning the integrated fusion model. Overall the study required approximately 5500 GPU-hours (∼7.6 GPU-months). We used a combination of H100 and A100 GPUs for the work, depending on availability.

### 4.5 Embeddings for Genetic Perturbation Modeling

In the context of genetic perturbation modeling, we embed perturbations by embedding the gene that is being targeted. Below we detail the approaches we consider.

#### 4.5.1 Expression

Unlike the other embedding categories, all of the expression-based gene embeddings are contextualized in the sense that they are based on control cell data from a given cell line.

##### Preliminaries: Fixed vs. variable gene set aggregation

The expression-based FMs described below are used to map a collection of single cells *X* ∈ ℝ^*N×G*^ to a collection of *d*-dimensional gene embeddings *Z* ∈ ℝ^*d×G*^. These single cells are typically the control cells for a particular cell line.

For some models (AIDO.Cell, scGPT, scPRINT) the gene set is the same for every cell. In this case, we obtain *Z*^*′*^ ∈ ℝ^*N×d×G*^ and simply compute

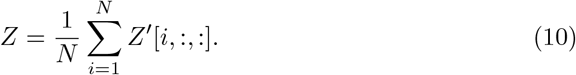

However, there are other models (Geneformer, TranscriptFormer) for which the gene set used to encode a cell depends on the cell itself. Let *Q*_*g*_ ⊂{1, …, *N}* be the set of cell indices for which gene *g* is used to encode the cell (and thus an embedding for *g* exists). As we iterate through cells, we obtain tuples (*g, z*_*i*_(*g*)) where *g* ∈ {1, …, *G*} denotes a gene and *z*_*i*_(*g*) denotes the embedding of gene *g* from cell *i*. Then we compute

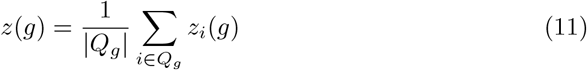

and form *Z* = concat([*z*(1), …, *z*(*G*)]). Thus, some gene embeddings may be more stably estimated than others depending on the size of *Q*_*g*_.

##### AIDO.Cell 3M/10M/100M [58]

AIDO.Cell is a family of transformer-based cellular foundation models, pretrained on 50 million cells spanning a wide range of tissues. AIDO.Cell performs dense self-attention over ∼18,000 protein-coding genes, producing a *d*-dimensional contextualized embedding for each gene for each cell. To compute gene embeddings for a cell line, we compute the average gene embeddings over all control cells. The gene set is fixed. We also consider the learned positional encodings from the AIDO.Cell (3M) model as embeddings (called “Positional” in e.g. Figure 1(a)) - this ablation allows us to measure the marginal benefit of the transformer layers.

##### Geneformer [28]

Trained on 30 million cells and adopts a language-model-inspired strategy. Instead of encoding all genes directly, it first ranks genes within each cell by expression and selects the top 2048. These “top-ranked genes” are treated as “tokens,” forming a cell-specific “sentence” that is processed by transformer layers. Given that this set of genes varies from cell-to-cell, we use the variable gene set aggregation method described above.

##### scGPT [30]

A transformer-based foundation model trained on 33 million cells that bins the expression values of genes before obtaining a gene-specific embedding per cell. The gene set is fixed.

##### scPRINT [59]

A cellular foundation model trained on 50 million cells designed to infer gene regulatory networks. It uses a bidirectional multi-head transformer and optimizes a zero-inflated negative binomial loss. The gene set is fixed.

##### TranscriptFormer [60]

A generative, multi-species foundation model trained on 112 million single cells from 12 species. It uses a fixed context length of 2048 “genes”, but this vocabulary varies depending on the species and context of the input. Given that this set of genes varies from cell-to-cell, we use the variable gene set aggregation method described above.

#### 4.5.2 Prior Knowledge

##### DepMap [61]

An embedding may capture gene function without being useful for predicting the effect of a gene knockout. For instance, consider a gene whose loss can be compensated for by another. To construct embeddings that capture this sort of information, we collect gene essentiality measurements for over 400 cell lines from the DepMap Portal. This yields a matrix *A*∈ ℝ^*N×G*^ where *N* is the number of cell lines and *G* is the number of genes for which essentiality scores are measured. To represent a gene, we simply use the corresponding column *z*_*k*_ = *A*[:, *τ* (*k*)] where *τ* (*k*) indexes the gene corresponding to perturbation *P*_*k*_.

##### GenePT [62]

GenePT produces gene embeddings by taking text descriptions of genes from the biomedical literature and using an LLM (GPT-3.5) to extract embeddings of that text. We consider 6 variants of embeddings from GenePT, one based on NCBI gene cards and UniProt gene descriptions and 5 based on gene ontology information.

##### GenotypeVAE [63]

GenotypeVAE learns gene embeddings by training a variational autoencoder to learn latent embeddings of gene ontology annotation vectors. The original GenotypeVAE was trained with a latent dimension of 10. For the embeddings in this work, we retrained the model with a latent dimension of 128 to increase the capacity of the model. Additionally, we used the latest gene annotations from Functionome [64] to encode genes. All hyperparameters were identical to those from the original GenotypeVAE.

##### STRING GNN

We train a simple graph convolution network with residual skip connections using the link prediction objective on StringDB [65], a protein-protein interaction network. Node embeddings are randomly initialized. We used around 300 Optuna trials to tune over layers (2, 4, 6, 8), dimensionality (128, 256), dropout (0, 0.1, 0.2), learning rate (1e-3, 1e-4), weight decay (0, 0.01), batch size (1024, 2048, 4096, 8192, 16384), StringDB graph filter threshold (0.6, 0.7, 0.8, 0.9), and layer type (Graph Isomorphism Network (GIN), Graph Convolutional Network (GCN)). Optimal values are underlined. We performed a 90:10 train to validation split of the StringDB edges and selected the model with the lowest validation loss on the link prediction objective.

##### STRING Network [66]

Network-based embeddings were obtained by applying weighted node2vec to species-specific STRING protein-protein association networks, resulting in 128-dimensional node representations.

##### STRING Spectral

Starting from StringDB, we construct a weighted interaction network using only the experimental, coexpression, and database channels. Interactions were filtered to maintain a medium-confidence threshold, retaining only those with a combined weight greater than 400. Using the `SpectralEmbedding` function from scikit-learn, the resulting adjacency matrix was projected into a 128-dimensional space. This spectral approach utilizes Laplacian eigenmaps to transform the graph’s topological structure into Euclidean vectors, ensuring that proteins with functional associations are positioned closely together in the embedding space.

##### WaveGC [67]

WaveGC is a wavelet-based graph convolutional network. We use 2 layers, 3 wavelet scales, and 7 Chebyshev terms. The model was trained using the Adam optimizer with a learning rate of 1e-3 for 200 epochs. We select the model with the best validation loss. We use early stopping with a tolerance of 30 epochs. The model was trained on link prediction using the STRING network.

#### 4.5.3 DNA

##### AIDO.DNA [34]

We use the AIDO.DNA model to compute gene embeddings based on the nucleotide sequence. For each gene, we run inference on a 4kbp window centered at the transcription start site. This yields per-nucleotide embeddings, which are then mean pooled to form gene embeddings.

#### 4.5.4 Protein

In all cases below, multiple protein isoforms may be available for a given gene. In such cases, the embedding for a gene is the mean over embeddings of corresponding isoforms. For models that produce per-residue embeddings, protein embeddings are produced by mean pooling over residues.

##### AIDO.StructureTokenizer [68]

300M parameter VQ-VAE model for protein structure tokenization.

##### AIDO.ProteinIF-16B [69]

16B parameter protein language model based on a sparse mixture of experts.

##### AIDO.Protein-RAG-16B [36]

A multimodal protein language model that integrates Multiple Sequence Alignment (MSA) and structural data.

##### ESM2 [35]

650M parameter protein language model trained on single protein sequences, without using MSA data.

##### STRING Sequence [70]

Protein sequence embeddings from the ProtT5, an encoder-decoder transformer model.

### 4.6 Embeddings for Chemical Perturbation Modeling

#### Molecular fingerprints

Fingerprints are fixed-length vector representations of molecules, obtained by hashing the presence or absence of specific chemical substructures (e.g., circular subgraphs, paths, atom environments). They are fast to compute and widely used in cheminformatics. We use several from RDKit [71]: MACCS, Topological, SECFP, ECFP:2, Avalon, and ErG.

#### SMILES-based foundation models

These models - MiniMol [72], Chem-BERTa [37], Uni-Mol [73], MolT5 [74] - are large pretrained transformers that operate on either a SMILES string or a molecular structure graph. These models are trained with different supervised and self-supervised objectives, including contrastive learning between text descriptions of the drug and structure-based embeddings of the drug (MoIT5), multi-target regression on chemistry-related labels from RDKit (ChemBERTa-MTR, MiniMol), masked atom prediction (Uni-Mol), and masked language modeling on the SMILES string (ChemBERTa-MLM),

#### Text embeddings

These embeddings (ChatGPT) use an off-the-shelf LLM (in our case, `text-embedding-3-small` from OpenAI) to embed textual metadata. In particular, we form a prompt that includes the name of the drug, the SMILES representation, any known targets, the mechanism of action (if known), and clinical trial status.

#### Target embeddings (FMs)

Many small molecules have known or suspected gene targets. To put this into practice for a given chemical perturbation dataset, we first use an LLM to predict 3 genetic targets for each small molecule given the name of the drug. (We make no claim about the accuracy of these targets - we simply evaluate their utility on the downstream task of perturbation response prediction.) We use `claude-4-sonnet-20250514`. We then use expression FMs (scPRINT, AIDO.Cell 100M, TranscriptFormer) to embed those genes. For simplicity, we use the embeddings derived from the Norman dataset. Using the control cells from the chemical perturbation dataset itself may perform better. The embeddings of the three targets are concatenated.

#### Target embeddings (ablation)

As an ablation on the FM-based target embeddings, we also experiment with encoding the targets as binary vectors (1 if a gene is a target, 0 otherwise; the number of genes is equal to the number of unique targets in the dataset). This method is denoted Targets Binary (Name). We also try a weighted version where we ask the LLM to return targets along with confidence weights: 1.0, 0.5, or 0.25. Then the target encoding replaces the “1” values with the corresponding weights. This method is named Targets Weighted (Name). Finally, we also try providing PubChem descriptions for each drug in addition to the drug name when querying for targets. This results in the methods Targets Binary (Name, PubChem) and Targets Weighted (Name, PubChem). We again use `claude-4-sonnet-20250514` for target prediction.

#### Affinity-based embeddings

Boltz-2 [75] is an open-source implementation of AlphaFold 3 that predicts 3D protein-ligand structures and estimates binding affinity. In this work, we used Boltz to develop embeddings of small molecules based on their predicted interactions with various proteins.

For each small molecule, we co-folded and computed the predicted binding affinity with a set of proteins. This allowed us to create a vector where each entry corresponds to the predicted binding affinity between the molecule and a given protein. We can treat this vector as an embedding of the small molecule.

We consider two variants: fragment-based embeddings and protein-based embeddings. See Supplemental Methods for details.

### 4.7 Perturbation Prediction Models

#### 4.7.1 Positive Controls

To establish what strong performance would look like, we considered several positive control methods.

##### Embedding oracle

Let *D* ∈ ℝ^*K×G*^ be the matrix of treatment effects for *G* genes over *K* perturbations. Crucially, let these *K* perturbations include both the train and test data. Choose an embedding dimension *d*. We perform PCA on *D*, resulting in *K* embeddings of dimension *d*.

Intuitively, these embeddings directly encode the similarity between all perturbations in the dataset, including the test set. Formally, the Eckhart-Young theorem [76] implies that these are the *d*-dimensional embeddings that compress *D* optimally (in terms of L2 error, and assuming a linear decoder). Note that nonlinear models could in theory produce *d*-dimensional embeddings that compress *D* with lower error, but even so we expect this baseline to be difficult to beat.

##### Noise oracle

To complement the Embedding Oracle, we introduce another oracle that uses bootstrapping to characterize the variability of the estimate of the treatment effect from the test data.

Let *X* ∈ ℝ^*N×G*^ be a matrix of expression measurements for *G* genes in *N* cells, some of which receive a perturbation and some of which are control cells. Given control and perturbed cells, we can compute the perturbation’s treatment effect which we will call Δ ∈ℝ^*G*^.

By sampling with replacement from *X* and repeating *M* times, we can create a collection of *M* bootstrap samples 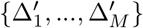. These samples can be transformed to samples of the error in estimating Δ by computing

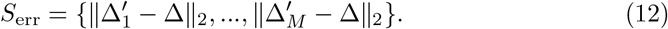

The samples in *S*_err_ approximate the distribution of the error in estimating Δ from repeated experiments. Then we define *ϵ*(*q*) - the noise oracle at *q* - to be the *q*th quantile of *S*_err_.

For example, if a model’s prediction 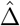 satisfies 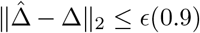, then the noise oracle is higher than the model’s error 10% of the time. We believe this is a challenging standard for any model to reach. Throughout this work we use *q* = 0.9. In the same vein, we can recompute DEGs under resampling to get a distribution of F1 scores and define a corresponding noise oracle for the DEG classification setting. In this case, since higher is better, we use *q* = 0.1.

Note that in practice we perform a hierarchical bootstrap, where we bootstrap over batches (*M*_outer_ trials) and cells within batches (*M*_inner_ trials) with the total number of trials being *M* = *M*_outer_ × *M*_inner_. See Supplemental Methods for details. We consider additional oracle formulations in Section A.3.5.

#### 4.7.2 Negative Controls

To establish null performance, we considered several negative control methods which we expect any strong method to beat.

##### Train Mean

For LFC regression, the Train Mean is a model that predicts the mean response over all training perturbations. For DEG classification, the Train Mean samples predictions from the distribution of classes observed for each gene in the training set.

##### No Change

For any perturbation *P*_*k*_, this method simply predicts that *P*_*k*_ has no effect. That is, we predict 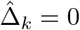 for all *k*, meaning that the perturbed expression was identical to control expression (in the LFC regression setting) or that no genes are differentially expressed (in the DEG classification setting).

##### Random Embeddings

For each perturbation *P*_*k*_ (gene or small molecule), we define a random embedding *z*_*k*_ as follows. Given an embedding dimension *d*, we sample for each perturbation an embedding *z*_*k*_ ∼ Uniform([0, 1]^*d*^). These embeddings carry no information about true similarity between perturbations and therefore cannot enable any generalization. Predictive models (e.g. kNN, Lasso) are fit using these embeddings under our standard benchmarking protocol.

##### PCA

A simple and natural way to produce gene embeddings is to perform PCA on the cells in the control population. Formally, let *X* ∈ ℝ^*N×G*^ be a matrix of expression values for *G* genes over *N* control cells for some cell line. Choose an embedding dimension *d*. Then by performing PCA on *X*^*T*^ we can produce *G* embeddings of dimension *d* = 128.

#### 4.7.3 Classical Machine Learning Methods

We tested several standard machine learning models including lasso, linear regression, and kNN regression. We detail our use of these models in Supplemental Methods, where we discuss embedding evaluation protocols.

#### 4.7.4 Advanced Perturbation Prediction Methods

In this section we describe the advanced perturbation prediction methods we evaluate in this work. For consistency, all models are evaluated using our LFC regression formulation, even those that make single-cell predictions (Latent Diffusion, Flow Matching, Schrödinger Bridge). For each perturbation, control cells are sampled within each batch and passed through the model conditioned on the perturbation embedding to generate predicted perturbed single-cell profiles. These predictions are averaged to form per-batch predicted pseudobulks, which are then averaged across batches. The final prediction for each perturbation is obtained by subtracting the global control pseudobulk from the batch-averaged predicted pseudobulk, resulting in predictions 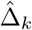 for each perturbation *P*_*k*_.

##### GEARS

GEARs [27] is a graph neural network where the edges are constructed roughly based on cell-ontology. Specifically, there is a cell-state encoder that is a GNN, and a perturbation encoder that is also a GNN. The training scheme is that one cell is sampled from the control distribution, and the two GNNs are used to predict the gene expression of the perturbed cell. This model is trained on one cell type at a time.

##### Latent Diffusion

Diffusion models are a powerful class of generative models that learn to iteratively transform a Gaussian random vector into a high quality data sample [45, 77]. (If we perform modeling in a latent space instead of data space, this is termed *latent* diffusion.) Applications to perturbation modeling include Squidiff [43] and dbDiffusion [78].

In general, diffusion models have a forward and reverse process. In the forward process, a data sample *x*_0_ has small amounts of noise progressively added until the data sample is fully transformed into a Gaussian random noise sample *x*_*T*_ after *T* steps. In the reverse or generative process, the model *P*_*θ*_(*x*_*t−*1_ | *x*_*t*_) learns to predict *x*_*t−*1_ from *x*_*t*_ by training a neural network (parameterized by *θ*) to predict the noise that was added to *x*_0_ to produce *x*_*t*_. At prediction time, the generative model is iteratively applied to new Gaussian noise samples to generate data samples. Generation is conditioned on an embedding of the perturbation being applied. See Supplemental Methods for implementation details.

##### Flow Matching

Flow Matching models can be seen as a generalization of diffusion models where, instead of transforming a sample from a Gaussian distribution to the data distribution, the sample can be transformed from one arbitrary distribution to another [46]. Applications in perturbation modeling include CellFlow [42] and CellFlux [79].

In Flow Matching, the model is trained to learn the vector field between two distributions by sampling linear interpolants during training. Specifically, for each training sample, *x*_0_ is sampled from the start distribution, *x*_1_ is sampled from the target distribution, and *t* ∈ [0, 1] is a sampled time point. A linear interpolant is drawn between the samples *x*_*t*_ = (1 − *t*) * *x*_0_ + *t* * *x*_1_ and a derivative is calculated at the point *dx*_*t*_*/dt* = *x*_1_ − *x*_0_. The model is trained with the mean squared error (MSE) objective to predict the velocity *dx*_*t*_*/dt* from *x*_*t*_ and *t*. At prediction time, Flow Matching uses ODE solvers to transform the start distribution into the target distribution according to the vector field learned by the model. Generation is conditioned on an embedding of the perturbation being applied. See Supplemental Methods for implementation details.

##### Schrödinger Bridge

Schrödinger Bridge models are a further generalization of diffusion and Flow Matching models that draws from the strengths of both [47]. This method has been applied to perturbation modeling in DEPARTURES [80] and ARTEMIS [44]. Schrödinger Bridge models allow for arbitrary distribution to arbitrary distributions (like Flow Matching) but are stochastic in nature (like diffusion). We adopt the implementation of simulation-free Schrödinger Bridge models from Tong et al. [81], which builds on the aforementioned diffusion and Flow Matching models. Generation is conditioned on an embedding of the perturbation being applied. See Supplemental Methods for implementation details.

##### STATE

STATE [49] is a large transformer-based model trained using 100M perturbed cells. The model was developed for cross-context prediction, i.e. predicting the effect of perturbations seen during training in cell lines not seen during training. STATE operates at the level of cell populations, consuming a set of control cells and a perturbation encoding and predicting a set of perturbed cells. Implementation details for our analysis of STATE can be found in Supplemental Methods.

### 4.8 Integrating FMs to Predict Perturbation Results

#### 4.8.1 Preliminaries

Let *E*_*k*_ ⊂ *E* denote the set of embedding sources that are valid for perturbation *P*_*k*_, where *E* = {1, …, *J*} is the set of all possible embedding sources. Let 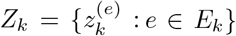 denote the set of valid embeddings for perturbation *P*_*k*_, where 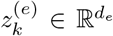. Typically *E*_*k*_ is a strict subset of *E*. (For example, in the genetic perturbation setting we might use a scRNA-seq FM as one gene embedding source *e*. But if the gene 22 corresponding to *P*_*k*_ was not used when training the FM, then 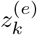 is not defined.) Let *L* be the total number of cell lines under consideration. Note that some embeddings are cell-line specific, so in general we should write *Z*_*k,l*_ to acknowledge this dependence. We omit this unless necessary for clarity.

#### 4.8.2 Simple Fusion Model

The aim of our embedding fusion model is to use the set of embeddings *Z*_*k*_ to predict the observed treatment effect Δ_*k*_ (for either the LFC regression or DEG classification case). We propose an attention-based fusion head for integrating embeddings, which we describe below. This model is trained jointly over multiple cell lines.

##### Architecture

In the first stage of our model, each embedding 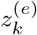 is passed through a preprocessing stage:

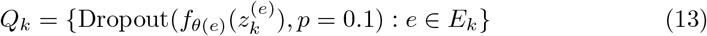

where 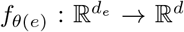 is a per-source projection model which maps embeddings of type *e* from dimension *d*_*e*_ to a common dimension *d*. We parameterize the cell line using a learnable, randomly initialized vector *z*_line_ ∈ ℝ^*d*^, which is preprocessed as

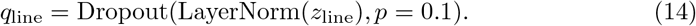

We also create a CLS token *q*_CLS_. We form our input set for perturbation *P*_*k*_ as

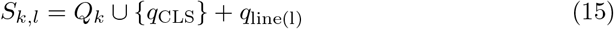

where addition is performed elementwise. Note that we do not explicitly inject source identity information into these tokens because the bias terms in the models *f*_*θ*(*e*)_ are able to perform that function.

Let *T*_*θ*_ : ℝ^*L×d*^ →ℝ^*L×d*^ denote a transformer model and let *g*_*ϕ*_ : ℝ^*d*^ →ℝ^*G*^ denote a prediction head. We integrate information across embeddings by computing *R* = *T*_*θ*_(*S*_*k,l*_). Finally, we make predictions as 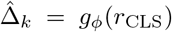 where *r*_CLS_ is the contextualized output (and thus a row of *R*) corresponding to *q*_CLS_.

##### Implementation details

We fix the transformer’s *d*_model_ = *d* = 100 and use 5 attention heads. We train with an L2 loss using a batch size of 64 and the Adam optimizer with a 5 epoch linear warmup. We use Optuna to tune over the depth of *T*_*θ*_ (ranging from 1 to 9 layers) and the learning rate (ranging from 1e-5 to 1e-2, searching logarithmically).

We also tune over a binary variable that toggles between the “global” cell line injection style presented above and an alternative approach where we append the cell line token to the input set. For the LFC regression setting on Essential (where fusion was quite successful), we found that the former consistently worked better than the latter.

We tune the simple fusion model with Optuna (TPE sampler) on the first fold only. Each trial trains on the first fold’s training perturbations minus a random 20% of perturbations for validation. Model selection is based on best-epoch validation loss (L2 for LFC, class-weighted cross-entropy for DEG). The test fold is never evaluated during tuning. The selected hyperparameters are then used to retrain all five folds.

##### Chemical perturbations

For sci-Plex and Tahoe we train only the simple fusion. It combines 10 (sci-Plex) or 29 (Tahoe) compound representations, the same for LFC regression and DEG classification. One model is trained per fold, jointly over all cell lines of the dataset. The cell-line embedding is added to every token, except for sciPlex DEG classification, where it is a separate token. For LFC regression, depth and learning rate were tuned with Optuna on the first fold, as above. The DEG models reuse hyperparameters chosen on earlier DEG labels and were not re-tuned on the variance-floored labels. On Tahoe, the LFC fusion converges to a near-constant prediction and is worse than the Train Mean.

#### 4.8.3 Full Fusion Model

This model is generally more complex and harder to tune, but it demonstrates that architectural innovations are available that can push the performance of multimodal fusion higher. This model may be trained per-cell-line or jointly over multiple cell lines. See Supplemental Methods for complete details.

## 5 Declarations

### Competing Interests

All authors are current or former employees of GenBio AI and may hold a financial interest in the company. GenBio AI provided the funding and resources for this research. EL performed this work as part of her consulting role for GenBio AI, not at Stanford University.

### Data and Code Availability

Data and code for this work can be found on GitHub: https://github.com/genbio-ai/foundation-models-perturbation

## A Supplemental Results

### A.1 Embedding Benchmarking

This section provides additional embedding benchmarking results for a variety of datasets.

#### A.1.1 Complete Results Table

A complete table of embedding benchmarking results is provided as a CSV in our GitHub repository. These values can be used to reproduce our figures.

#### A.1.2 Results for the Norman Dataset

We show LFC regression results using kNN in Figure A1. We observe that all methods perform poorly, with no method significantly outperforming Train Mean. There is a gap between the negative controls and the positive controls, indicating that better performance is possible in principle.

One possible issue is the small size of the Norman dataset. Prior work like Wu et al. [31] uses Norman for dual perturbation prediction. Our work considers only single perturbation prediction. This means we have 105 perturbations (before splitting), which may explain the poor performance across the board. It is also possible that the embeddings we consider are not well-suited for predicting the effects of CRISPRa. Without the availability of a large CRISPRa dataset it is difficult to test this hypothesis.

#### A.1.3 Results for the Essential Dataset

To complement the Lasso results in Figure 1, we show kNN results and rankings (Figure A2). We also show results for the DEG classification task (Figure A3). In all cases, the embedding rankings are largely similar to what we observe for LFC regression with Lasso (Figure 1) and kNN regression (Figure A2). For DEG prediction, the fusion models improve on Train Mean but not on the best single embeddings they are fusing.

#### A.1.4 Results for the Tahoe Dataset

We show LFC regression results for kNN (Figure A4(a)) and Lasso (Figure A4(b)). We also show results for the DEG classification task in Figure A4(c).

#### A.1.5 Results for the Sciplex Dataset

We show LFC regression results for kNN (Figure A5(a)) and Lasso (Figure A5(b)). We also show results for the DEG classification task in Figure A5(c).

### A.2 Evaluation of Advanced Methods

#### A.2.1 Advanced Perturbation Prediction Models

Figure A6 compares a strong unimodal embedding (WaveGC) against advanced methods like GEARS and three single-cell generative models (Latent Diffusion, Flow Matching, Schrödinger Bridge) conditioned on the same embedding. None of them convincingly improves on kNN.

**Fig. A1.**
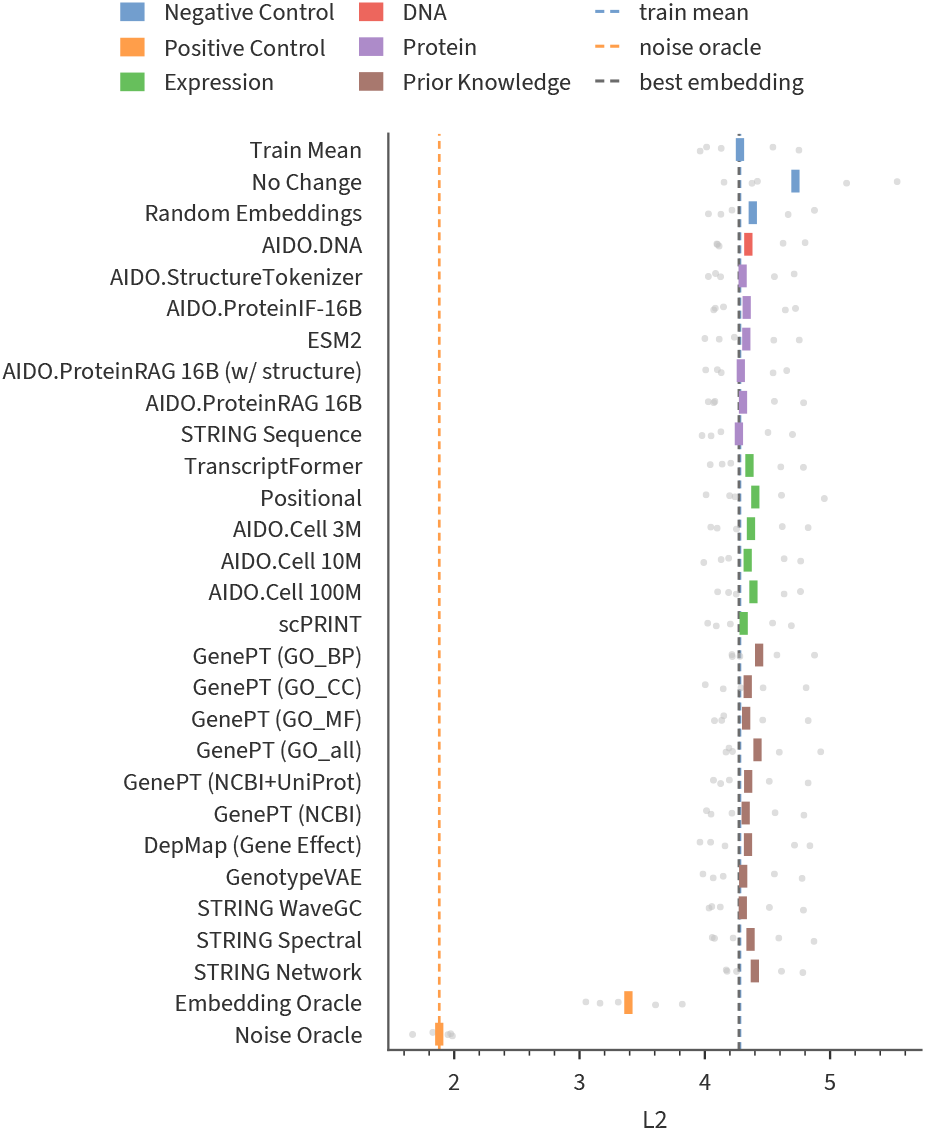
On the Norman screen, no gene embedding improves on the train mean. Out-of-fold LFC prediction with kNN on frozen gene embeddings for the single-gene CRISPR-activation perturbations of the Norman screen (K-562). L2 per method (lower is better), with rows grouped by family. Gray dots are folds and the colored tick is the mean over folds. Dashed lines mark the train mean, the noise oracle, and the best embedding. The train-mean and best-embedding lines almost coincide.

These runs used an earlier version of the Essential benchmark with folds drawn per cell line, so their L2 values are not comparable with the other Essential figures (caption of Supplementary Figure A6 and Supplementary Section B, Advanced Perturbation Prediction Method Details).

**Fig. A2.**
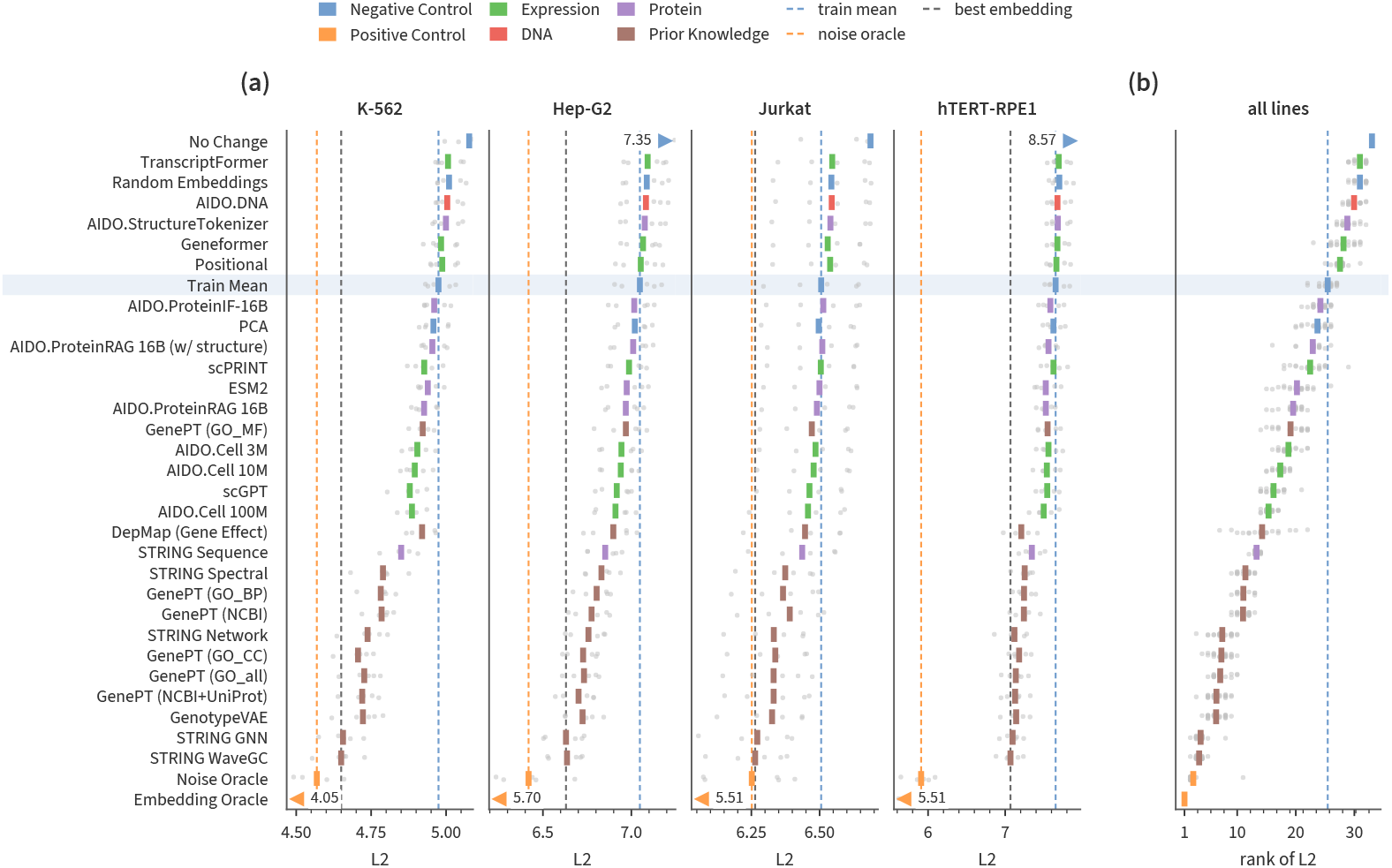
kNN regression mirrors the patterns of lasso on the Essential screens. Out-of-fold LFC prediction with kNN on frozen gene embeddings in the four Essential screens. (a) L2 per method and cell line (lower is better). Gray dots are folds and the colored tick is the mean over folds. Dashed lines mark the train mean, the noise oracle, and the best embedding. Means off the axis are drawn as triangles with their values. (b) Rank of L2 in each cell line and fold (Supplementary Section B.1). Rows left of the dashed line beat the train mean on average. Rows are shared by (a) and (b) and sorted by mean rank, best at the bottom.

**Fig. A3.**
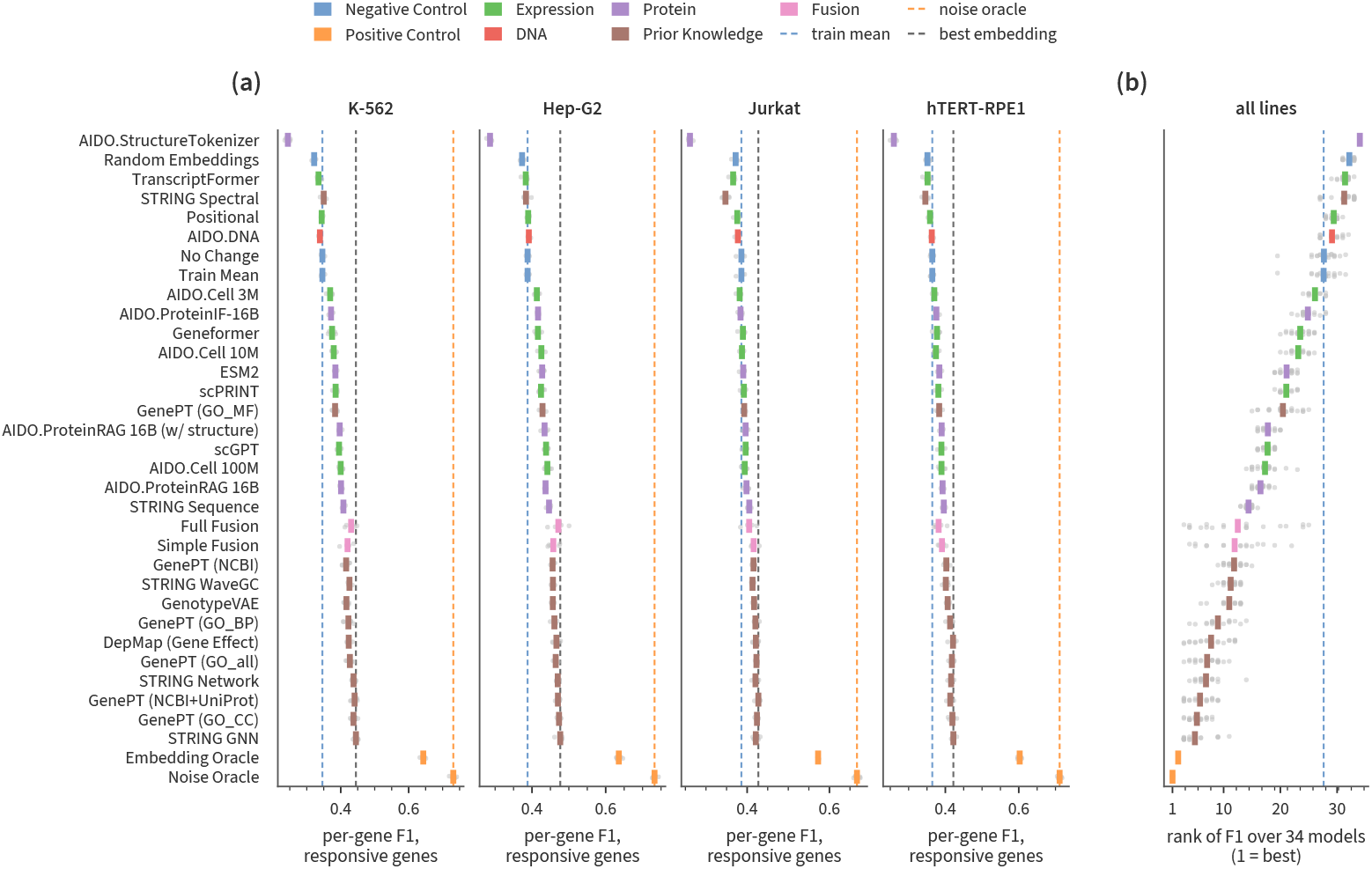
For DEG prediction, family rankings are similar to LFC but fusion does not help. Prediction of DE calls (down, not DE, up) from frozen gene embeddings in the four Essential screens. (a) Per-gene F1 over responsive genes per method and cell line (higher is better). Gray dots are folds and the colored tick is the mean over folds. Dashed lines mark the train mean, the noise oracle, and the best embedding. (b) Rank of F1 in each cell line and fold (Supplementary Section B.1). Rows are sorted by mean rank, best at the bottom.

**Fig. A4.**
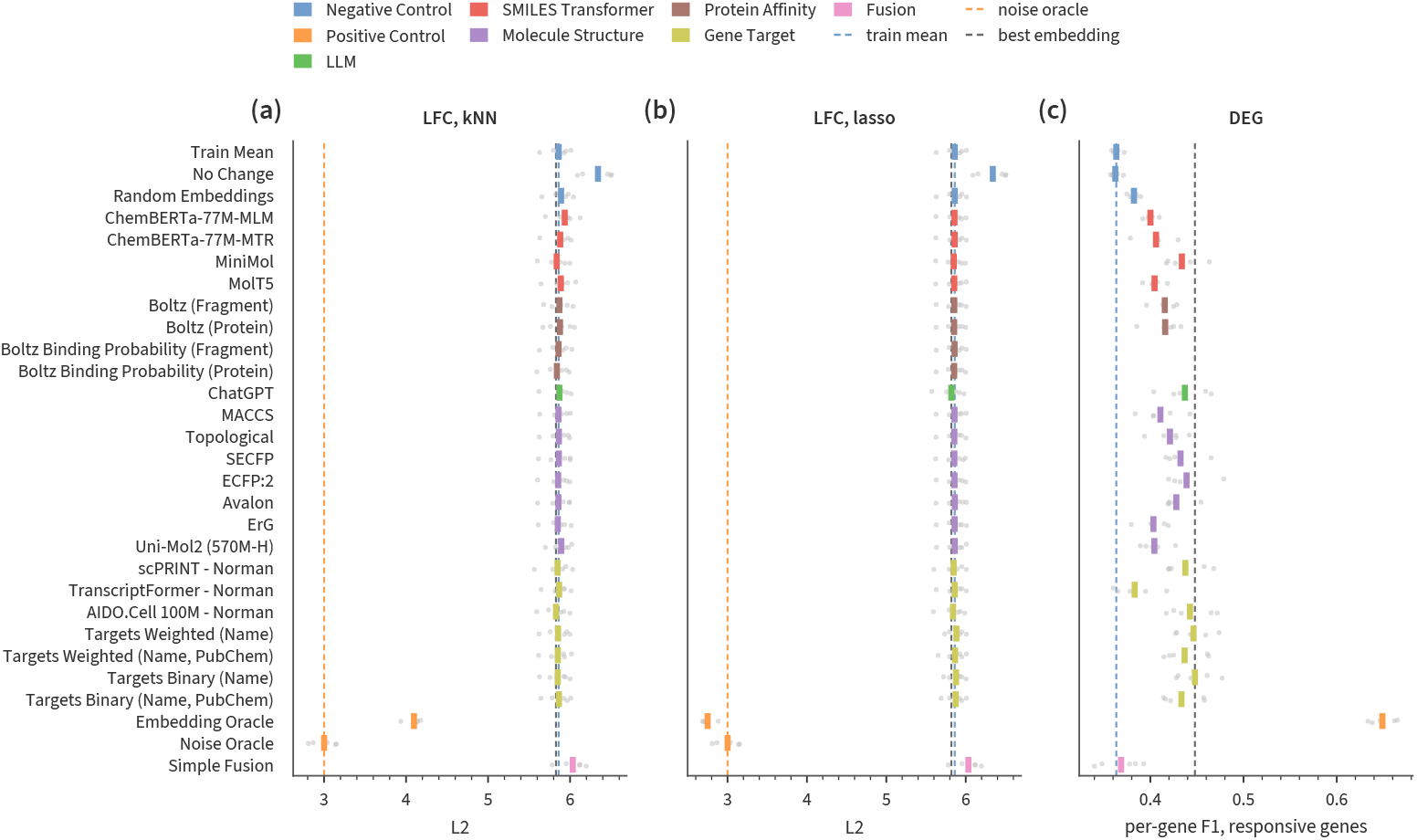
On Tahoe no compound embedding improves on the train mean for LFC, although several do for DEG prediction. Results on Tahoe, averaged over cell lines in each fold. (a, b) LFC prediction with kNN and with lasso, as L2 (lower is better). (c) DEG prediction, as per-gene F1 over responsive genes (higher is better). Rows are compound representations grouped by family, then controls, oracles, and the simple fusion. An empty row is a method not run on that task. Gray dots are folds and the colored tick is the mean over folds. Dashed lines mark the train mean, the noise oracle, and the best embedding.

**Fig. A5.**
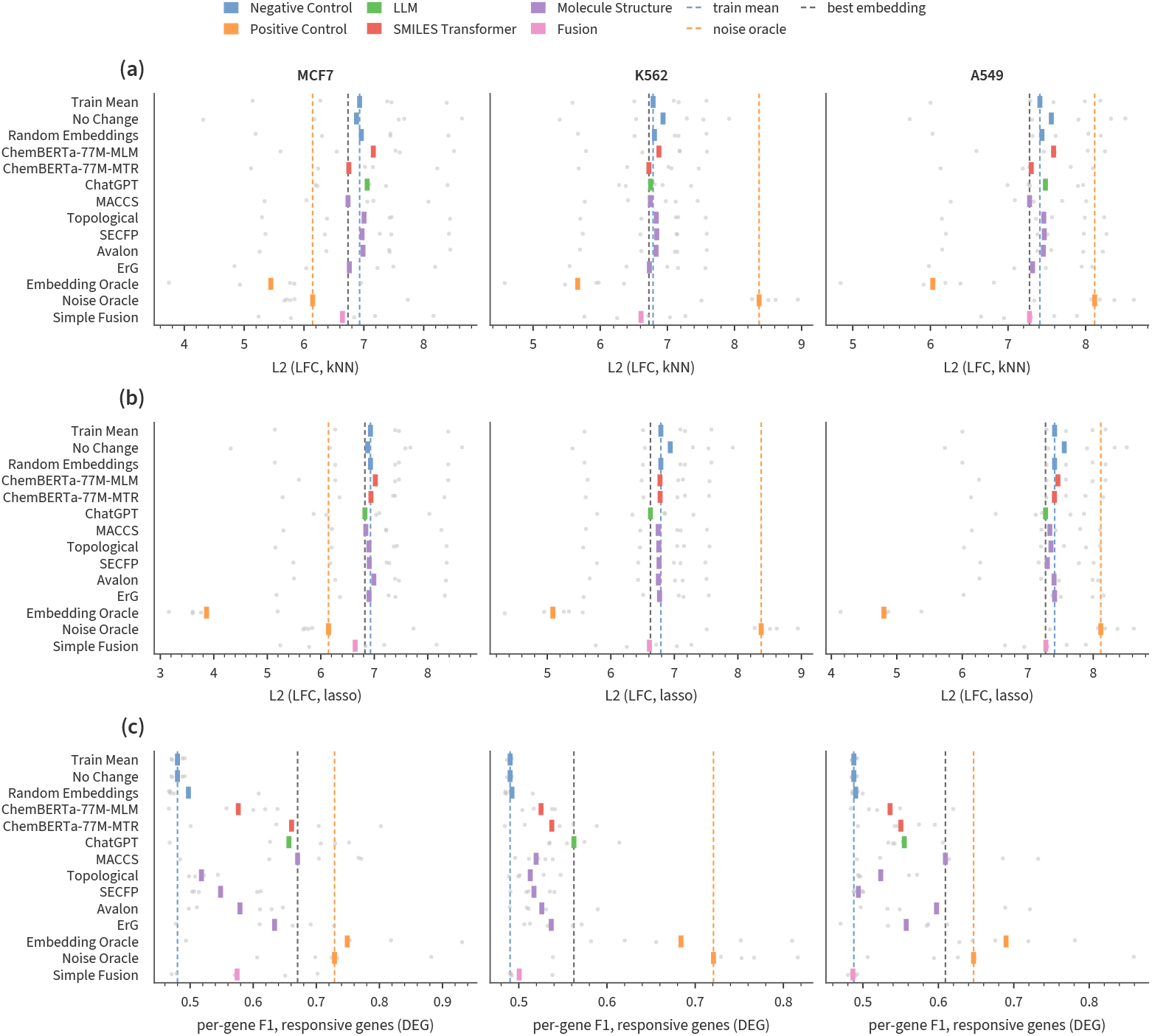
On sci-Plex, single compound embeddings marginally improve on the train mean for LFC, but fusion helps and several embeddings work for DEG prediction. Results on sci-Plex for MCF7, K562, and A549. (a, b) LFC prediction with kNN and with lasso, as L2 (lower is better). (c) DEG prediction, as per-gene F1 over responsive genes (higher is better). Rows are compound representations grouped by family, then controls, oracles, and the simple fusion. Gray dots are folds and the colored tick is their mean. Dashed lines mark the train mean, the noise oracle, and the best embedding. The simple fusion has its own head, so (a) and (b) show the same values.

**Fig. A6.**
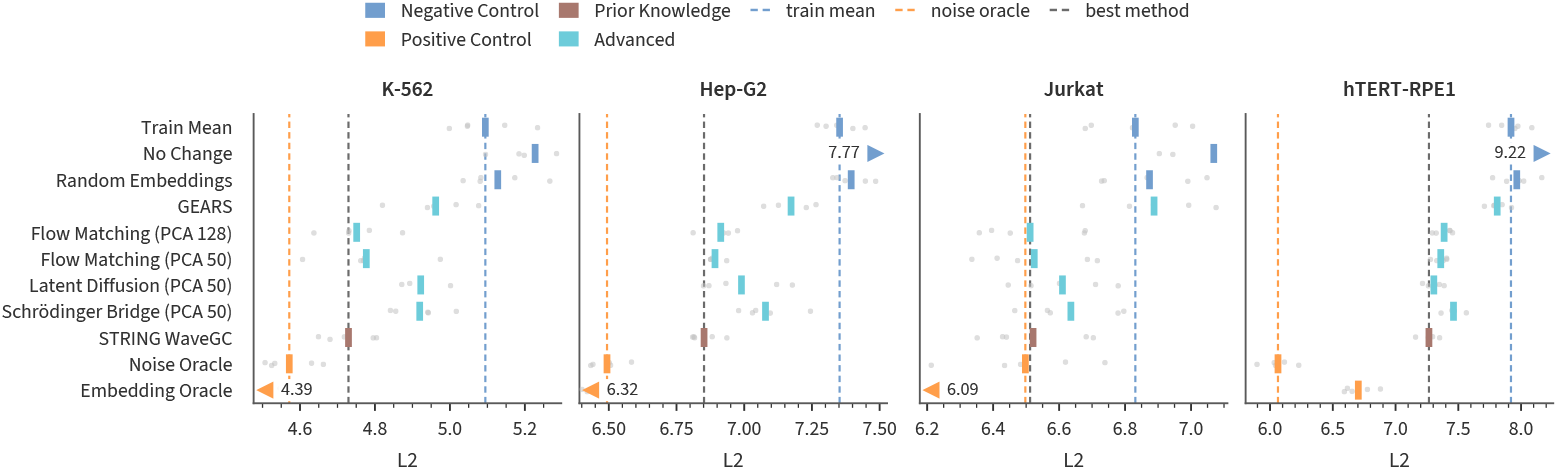
Generative and graph-based perturbation models do not improve on kNN with STRING WaveGC embeddings. L2 (lower is better) on the four Essential screens for GEARS, flow matching, latent diffusion, and a Schrödinger bridge, against kNN on STRING WaveGC, controls, and oracles. These methods were evaluated on an earlier version of the benchmark, with folds drawn per cell line. All references were recomputed on the same folds, so values are not comparable with Supplementary Figure A2. For latent diffusion and the Schrödinger bridge, the lower-test-error run of two is shown, which favors these methods. Gray dots are folds and the colored tick is the mean over folds. Dashed lines mark the train mean, the noise oracle, and the best non-oracle method. Means off the axis are drawn as triangles with their values.

**Fig. A7.**
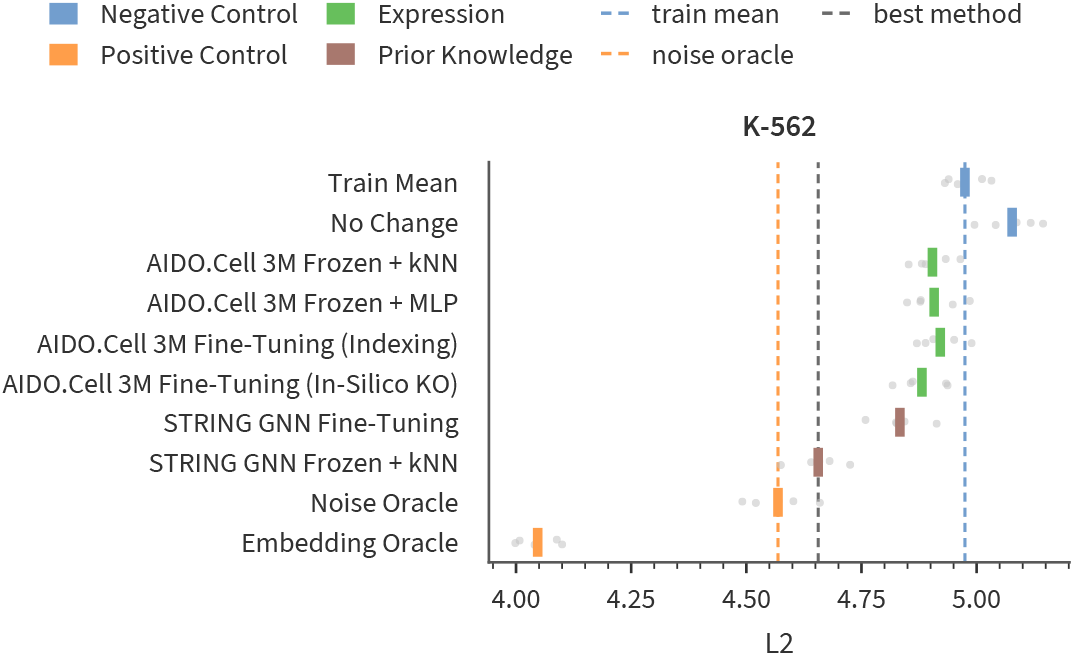
Fine-tuning does not improve on frozen STRING GNN embeddings with a kNN regressor on Essential K-562. L2 (lower is better) on Essential K-562 for frozen and fine-tuned AIDO.Cell 3M and STRING GNN, with controls and oracles. Gray dots are folds and the colored tick is their mean. Dashed lines mark the train mean, the noise oracle, and the best method. Fine-tuning variants are described in Section B.4.

#### A.2.2 Fine-Tuning

A comprehensive exploration of fine-tuning is beyond the scope of this work. However, we present a brief exploration in which we attempt to fine-tune two models (AIDO.Cell (3M) and STRING GNN) on the LFC regression task on Essential (Figure A7).

Our results are mixed, and indicate that fine-tuning is not a quick and easy win. The success of fine-tuning in our experiments depended on both the architecture and the fine-tuning method employed. For AIDO.Cell (3M), the In-Silico KO method (Supplemental Methods) did improve performance. This suggests that the model may be capturing task-relevant biological signals beyond simple embedding proximity. Conversely, the Indexing method for AIDO.Cell and our fine-tuning attempts for STRING GNN resulted in performance degradation relative to frozen embeddings. Overall, our fine-tuning results are mixed. More work is required to determine the most useful fine-tuning procedures for perturbation prediction.

#### A.2.3 Cross-Context Prediction

The primary focus of this paper is predicting the effect of unseen perturbations. In this section, we briefly consider the problem of predicting seen perturbations in unseen contexts. We study this cross-context formulation by using tasks from the STATE paper [49]. We show results for Essential (Figure A8, top) and Tahoe (Figure A8, bottom). We compare STATE against a kNN regressor and two MLP baselines (one on an embedding, one on one-hot encodings of perturbation identity). STATE is the best performer in terms of discrimination score on both Essential and Tahoe, but does not convincingly outperform the simple baselines in the other four cases we consider. On the Essential dataset, STATE performs at the same level as Train Mean.

**Fig. A8.**
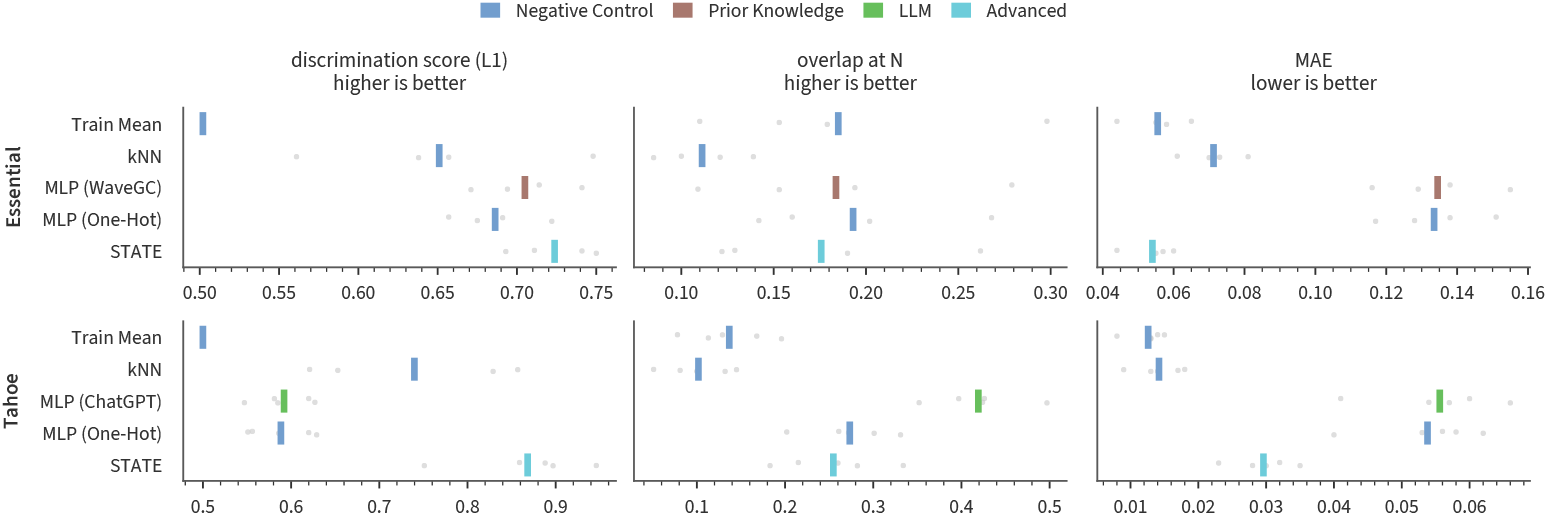
In cross-context prediction, STATE has the highest discrimination score on both datasets, while baselines match or beat it in two other metrics. Comparison with STATE on Essential (top) and Tahoe (bottom), scored with STATE’s evaluator, cell-eval (Supplementary Section B.1). Columns are the L1 discrimination score, DE-gene overlap at N, and mean absolute error (MAE). Rows are the train mean, kNN, an MLP on an embedding (STRING WaveGC for Essential, ChatGPT for Tahoe), an MLP on one-hot identities as an ablation, and STATE. Gray dots are held-out cell lines and the tick is the mean. Overlap at N uses the DE calls from cell-eval, not the variance-floored calls used elsewhere in this work.

### A.3 Additional Results

#### A.3.1 Genetic perturbation results replicate in the setting of Ahlmann-Eltze et al

To show that our findings are not an artifact of how we set up the task, we reevaluate our embeddings in the formulation of Ahlmann-Eltze et al. [32] using their exact code, splits, and metrics. Figure A9 shows our embeddings and the published results of Ahlmann-Eltze et al. [32] on the Replogle cell lines. We first note that many of our embeddings outperform all methods in that work. Second, we observe that the ordering of embedding families is largely the same as in our own evaluations: prior knowledge embeddings perform best, followed by an assortment of expression and protein embeddings, and finally DNA.

#### A.3.2 Embedding ranking is similar across datasets when sufficient signal exists

Our genetic perturbation results in the main paper focus on the Essential dataset. In this section we expand our results to perturbation screens from Nourreddine et al. [55] and Huang et al. [56] covering three additional cell lines. Figure A10 shows results for K-562 from the Essential dataset alongside these new cell lines. Results are stratified by per-perturbation SNR. When SNR is low, all methods typically collapse to the train mean. As SNR rises, we see the same pattern that we observed in the Essential dataset: prior knowledge performs best, followed by expression and protein foundation embeddings, followed by DNA embeddings. In almost all cases, all embeddings perform better with increasing SNR. In addition, Figure A11 shows that out fusion results also extend to the iPSC dataset for perturbations with sufficiently high SNR.

**Fig. A9.**
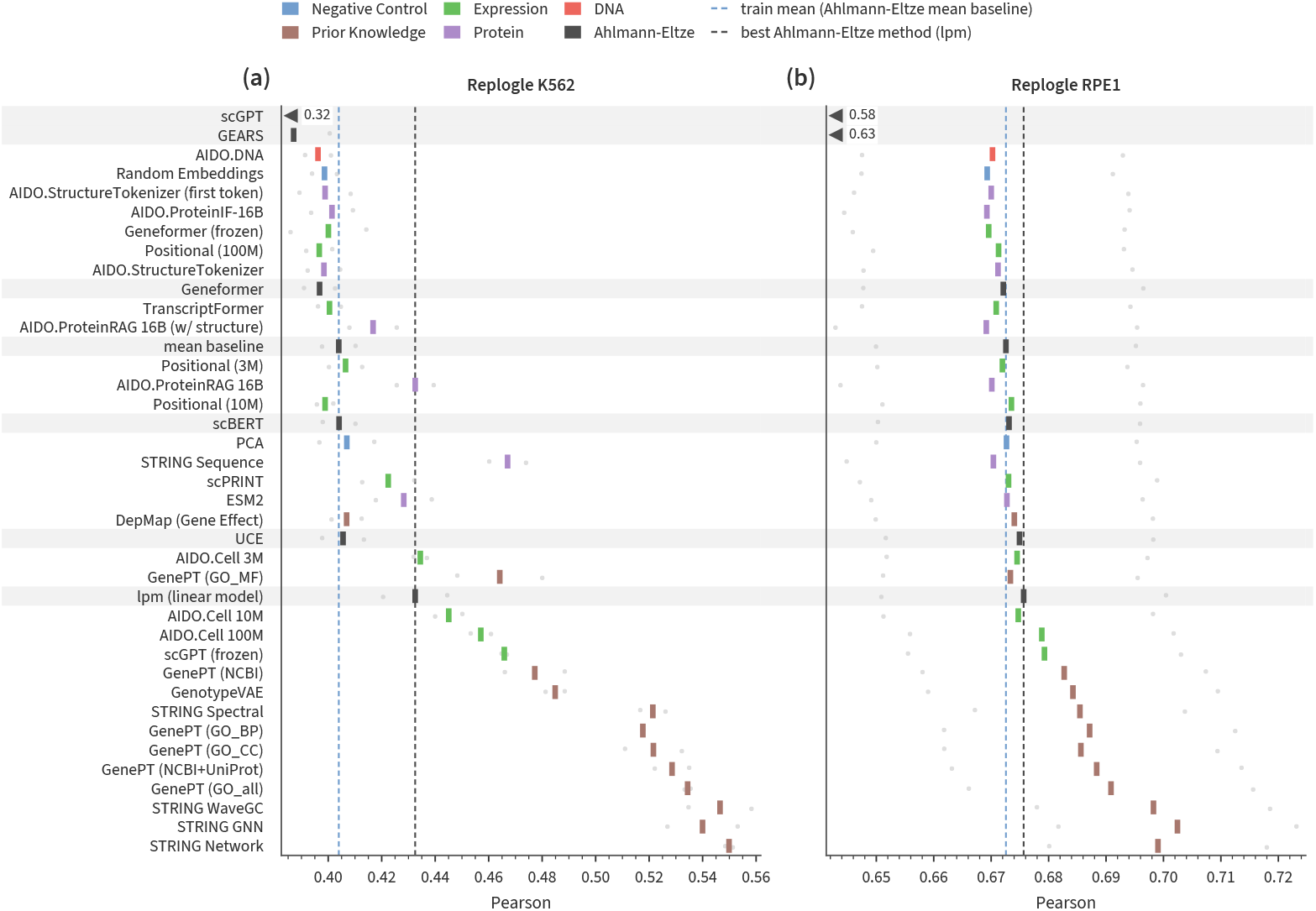
In the benchmark of Ahlmann-Eltze et al., some prior-knowledge and expression embeddings beat every published method. Pearson correlation between predicted and observed perturbation effects in the Replogle K562 (a) and RPE1 (b) screens. We use the code, splits, and metric of Ahlmann-Eltze et al. [32]. Our rows are kNN on frozen gene embeddings. Rows on a gray band are the published methods from that work. Gray dots are the two data splits, and the colored tick is their mean. Dashed lines mark the train mean (their mean baseline) and their best method (lpm). Means off the axis are drawn as triangles with their values. Rows are sorted by mean rank over both screens, best at the bottom.

It is interesting to note that different datasets have very different distributions of perturbations over SNR bins. The Essential dataset has mostly high-SNR perturbations, while the iPSC dataset shows the opposite pattern. The perturbations in the two cell lines from Xaira have a reasonably uniform distribution of SNR levels. This distribution determines what patterns are visible when one averages across all perturbations (e.g. dashed lines in Figure A10).

#### A.3.3 Embedding ranking is robust to more difficult splits

Unless otherwise noted, the results presented in this paper are based on IID random assignments of perturbations to train, validation, and test splits. It is therefore possible for highly similar perturbations to be assigned to different folds, which would likely make the task easier.

To make the problem more challenging, we can consider an alternative setting in which we hold out entire families of perturbations. We first map each gene to its most specific HGNC gene family, i.e. the deepest one in the HGNC family directed acyclic graph. In the case of a tie, a gene is assigned to the group with the lowest ID. Genes that cannot be assigned to a group through this procedure become their own singleton groups. For example, 84% of genes in Essential are assigned to proper non-singleton groups. Note that we do not include “host groups” which are defined by the presence of an intronic ncRNA as opposed to some sort of homology. Starting with the largest group, we create five folds by greedily assign each group to the fold with the fewest genes until all genes are assigned. To ensure comparability, we pool test results over folds before scoring. This means that, regardless of the fold construction policy, the test set will cover all perturbations.

**Fig. A10.**
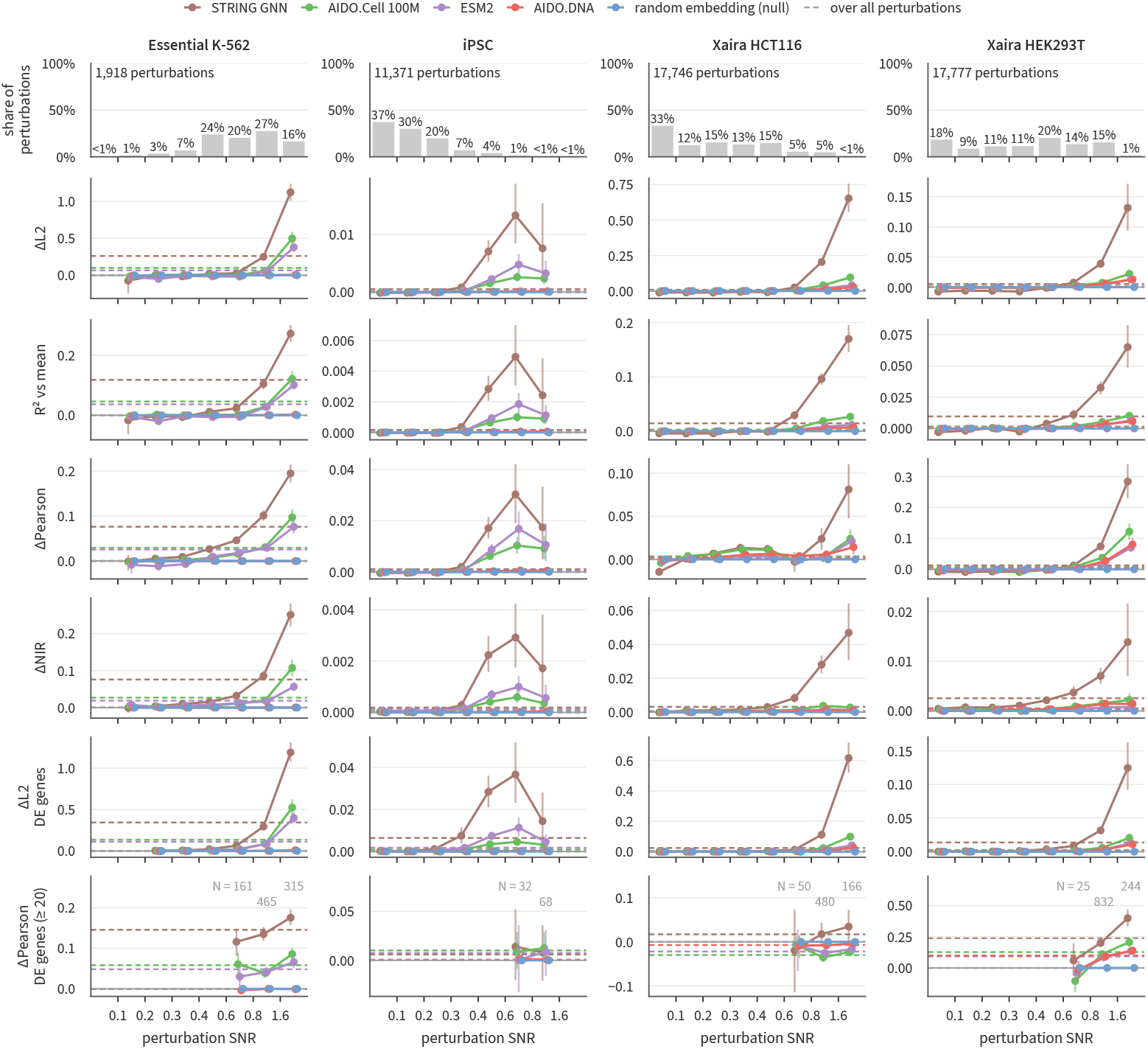
Embeddings improve on the train mean only for perturbations with replicable signal, and the share of such perturbations differs widely between datasets. Gain over the train mean, stratified by perturbation SNR, in Essential K-562, iPSC (KOLF2.1J), Xaira HCT116, and Xaira HEK293T. The top row is the share of perturbations per SNR bin. The rows below are ΔL2, R^2^ vs mean, ΔPearson, and ΔNIR on all genes, then ΔL2 and ΔPearson on DE genes (Supplementary Section B.1). Lines are one embedding per family and a random-embedding null. AIDO.Cell 100M embeddings are derived from each dataset’s own control cells. Dashed lines give each statistic over all perturbations. Gray numbers count the perturbations per bin in DE ΔPearson. Error bars are 95% bootstrap CIs over perturbations. All results use a lasso head.

**Fig. A11.**
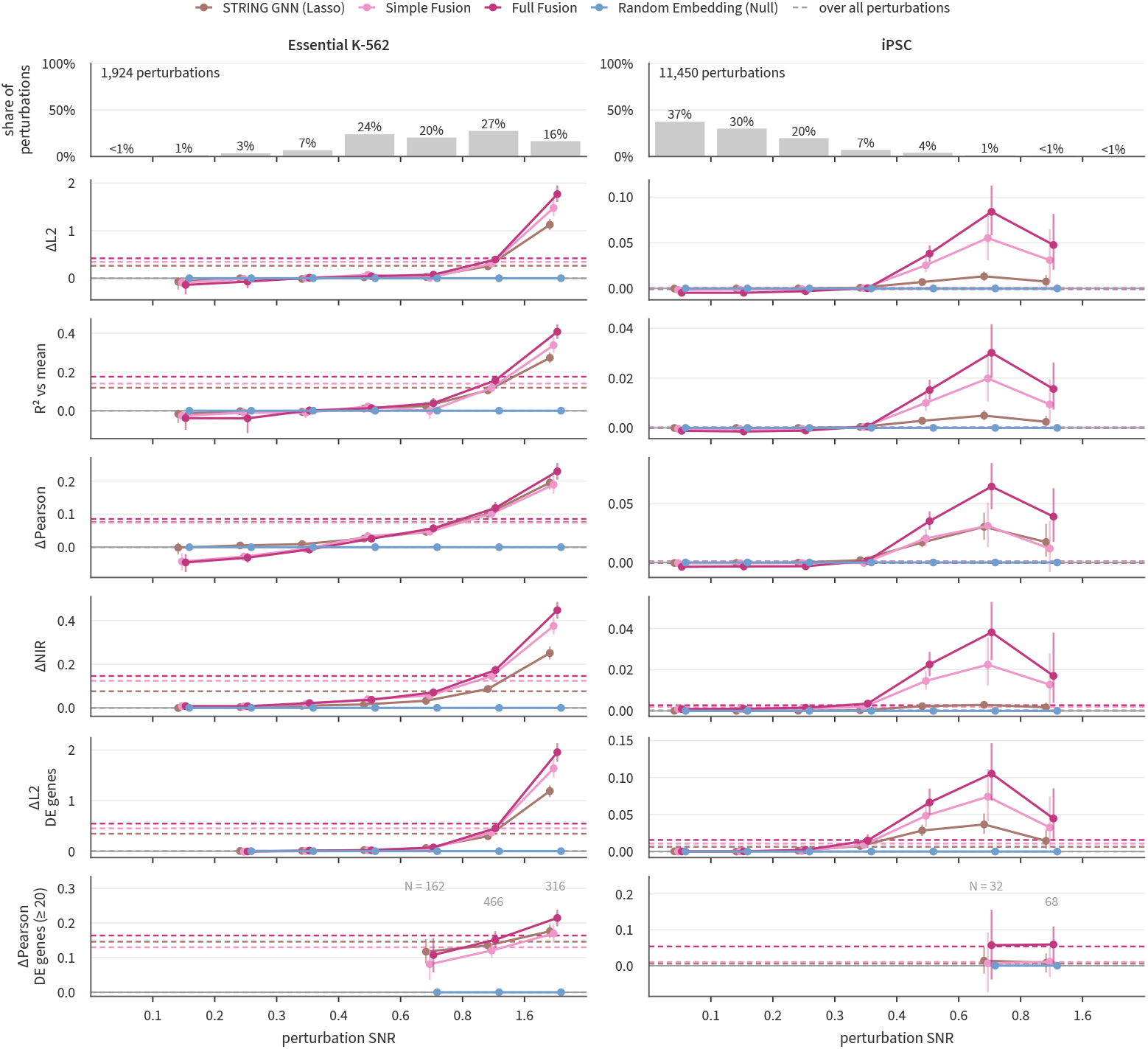
Gains from fusion sit at high SNR, like those of single embeddings. Gain over the train mean, stratified by perturbation SNR, for fusion models on Essential K-562 and iPSC (KOLF2.1J). The top row is the share of perturbations per SNR bin, among those every model scores. The rows below are ΔL2, R^2^ vs mean, ΔPearson, and ΔNIR on all genes, then ΔL2 and ΔPearson on DE genes (Supplementary Section B.1). Lines are lasso on STRING GNN, the simple and full fusion models, and lasso on a random embedding (null). On K-562 the fusions are trained jointly on the four Essential lines. On iPSC they are trained on iPSC alone. Dashed lines, gray numbers, and error bars are as in Supplementary Figure A10.

We build these family-level folds for two datasets (Essential K-562 and iPSC) and show the results in Figure A12. We observe two strong, consistent trends. First, the gene family splits are almost always more difficult than the IID splits. This can be observed by the downward slopes of the lines. Second, the method family ordering is nearly always: prior knowledge, expression, protein, DNA. The more challenging splits reduce performance, but do not change this basic pattern.

**Fig. A12.**
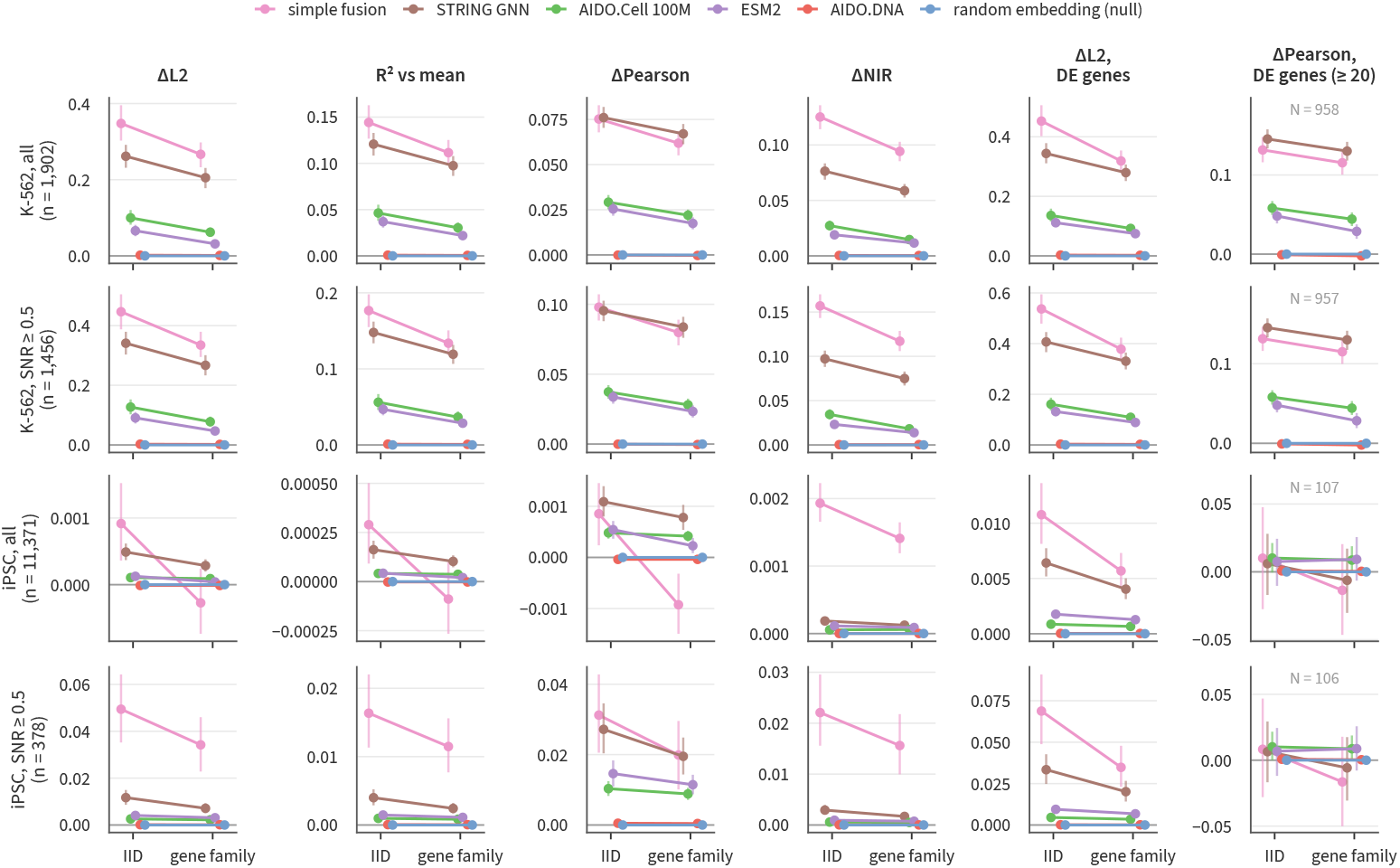
Holding out whole gene families lowers the gains but leaves the order of embeddings unchanged. Gains over the train mean with lasso under the IID split and under a split that holds out whole HGNC gene families, in Essential K-562 and iPSC (KOLF2.1J). Rows are each dataset over all perturbations and at SNR ≥ 0.5. Columns are ΔL2, *R*^2^ vs mean, ΔPearson, and ΔNIR on all genes, then ΔL2 and ΔPearson on DE genes (Supplementary Section B.1). A line joins each model’s values under the two splits, so lines that do not cross keep their order. Models are the simple fusion, one embedding per family, and a random-embedding null. The simple fusion was not re-tuned for the gene-family split. Error bars are 95% bootstrap CIs over perturbations.

#### A.3.4 Embedding rankings are stable across metrics and gene sets

There is not yet a consensus on which metrics are best suited for evaluating the accuracy of perturbation response predictions. In addition, there are many choices for which gene set to compute our metrics over (e.g. all genes, highly variable genes, differentially expressed genes). Figure A13 demonstrates that the rankings of methods are highly stable across gene sets and metrics. For the Essential dataset with all perturbations, the median agreement is 0.95 Spearman, with a minimum agreement of 0.91 Spearman. Method ranking is less consistent for the iPSC dataset if we use all perturbations, but strong consistency returns when we consider only high-SNR perturbations.

**Fig. A13.**
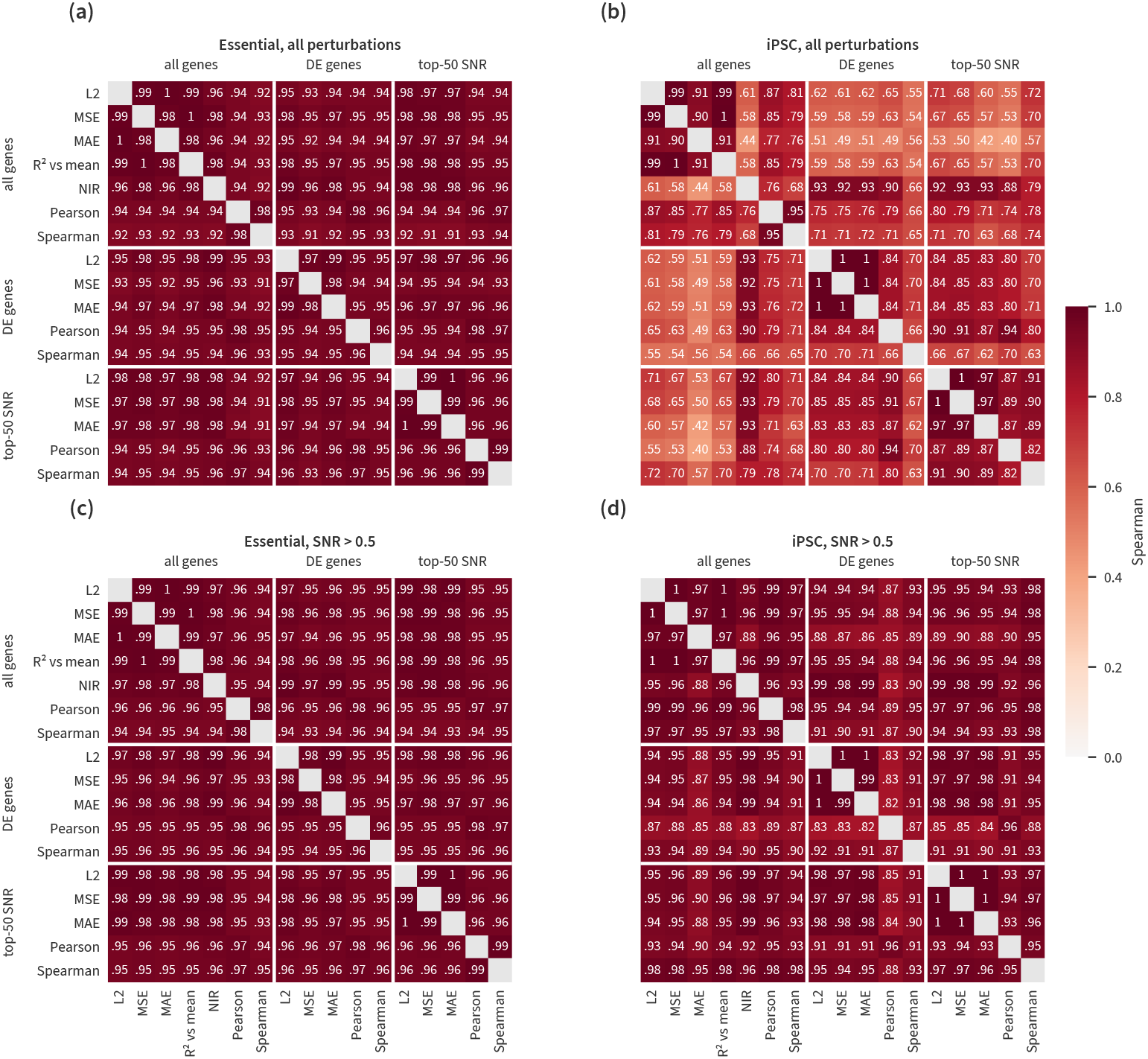
Metrics agree on the ranking of methods whenever the perturbations carry signal. Spearman correlation between method rankings for each pair of metrics, on all genes, DE genes, and top-50 replicate-SNR genes (Supplementary Section B.1). (a) Essential and (b) iPSC (KOLF2.1J), all perturbations. (c, d) The same, restricted to perturbations with replicate SNR *>* 0.5. Correlations are averaged over cell lines and the kNN and lasso heads.

#### A.3.5 Positive controls suggest different levels of headroom

It is useful to understand how much of the error in a perturbation prediction problem is irreducible and due to properties of the data itself. In the main paper, we take a step towards quantifying this irreducible error using the noise oracle and the embedding oracle (Section 4.7.1). In this section we construct additional variants of the noise oracle in an effort to shed more light on the different sources of irreducible error. For clarity we will call the noise oracle from the main paper the *bootstrap* noise oracle in what follows. We note that complementary work from Miller et al. [33] also highlights the importance for positive and negative controls for contextualizing performance numbers.

**Fig. A14.**
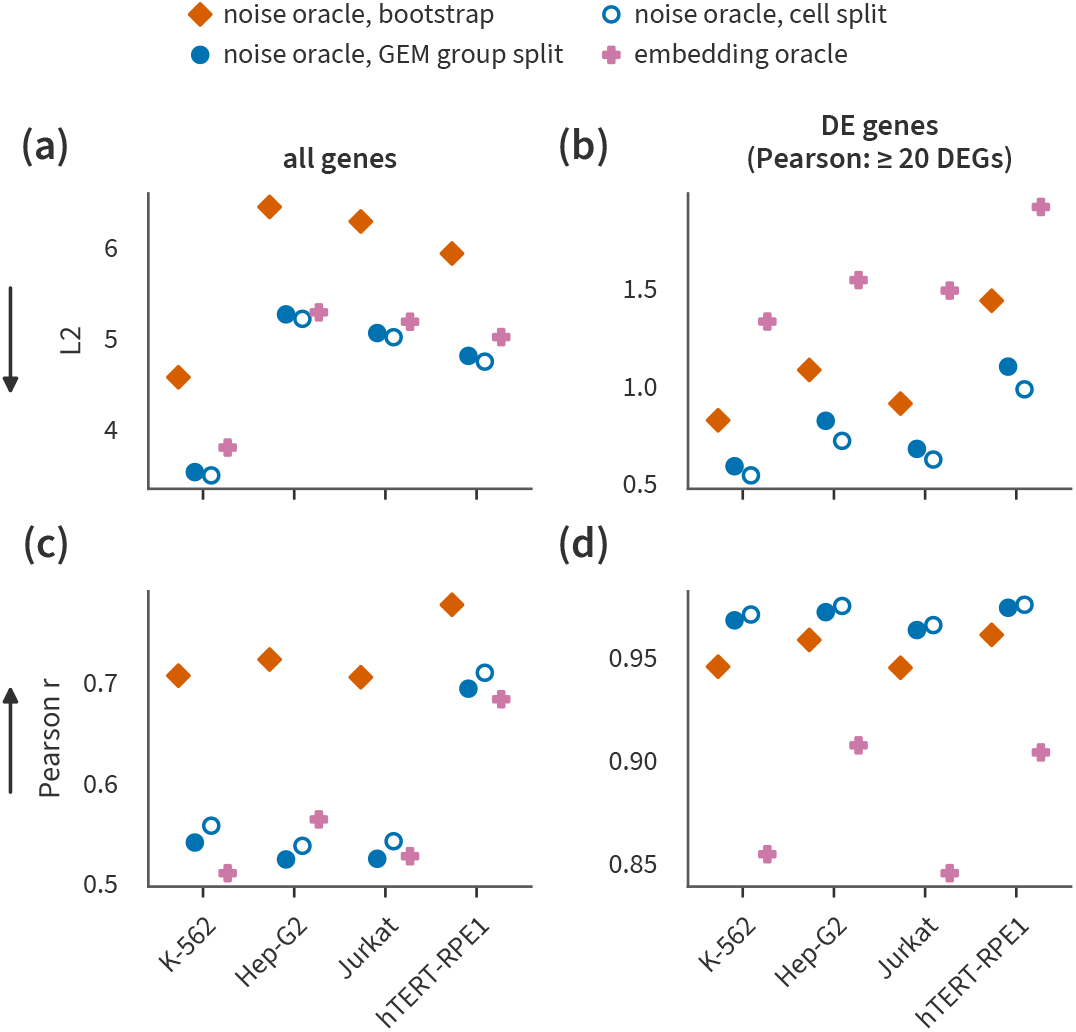
Different noise oracles disagree. Oracle references for Essential LFC prediction per cell line. Rows are L2 and Pearson *r*, with arrows marking the better direction. Columns are all genes (a, c) and DE genes (b, d). The noise oracle is estimated by the bootstrap and by two half-splits (Section B.10.2). The embedding oracle is drawn for comparison. Values are means over the perturbations that every source covers (Supplementary Section B.1).

Here we introduce two half-split noise oracles, which split each perturbation’s cells (cell split) or the GEM groups (GEM split) into two random halves and score the agreement between the two estimates (Section B.10.2).

Figure A14 compares the two oracles from the main text against the two replicate-based oracles. We note that the embedding oracle (pink) is more or less tied with the replicate-based oracles (blue) when we consider all genes (either L2 or Pearson). This means that 100 dimensions and linear projections suffice to capture almost all of the reproducible signal. The situation is different when we consider DE genes (L2 and Pearson), in which case there is a considerable gap between the replicate-based oracles and the embedding oracle. In that case, performance seems to be limited by the representation. Next, we note that the “worst” oracle (i.e. the one farthest from perfect performance) depends on the metric. However, in almost all cases it is either the bootstrap or the embedding oracle. Finally, the GEM group split and the cell split are almost always quite close. This suggests that the batch effects are small relative to variance within a perturbation for this dataset.

**Fig. A15.**
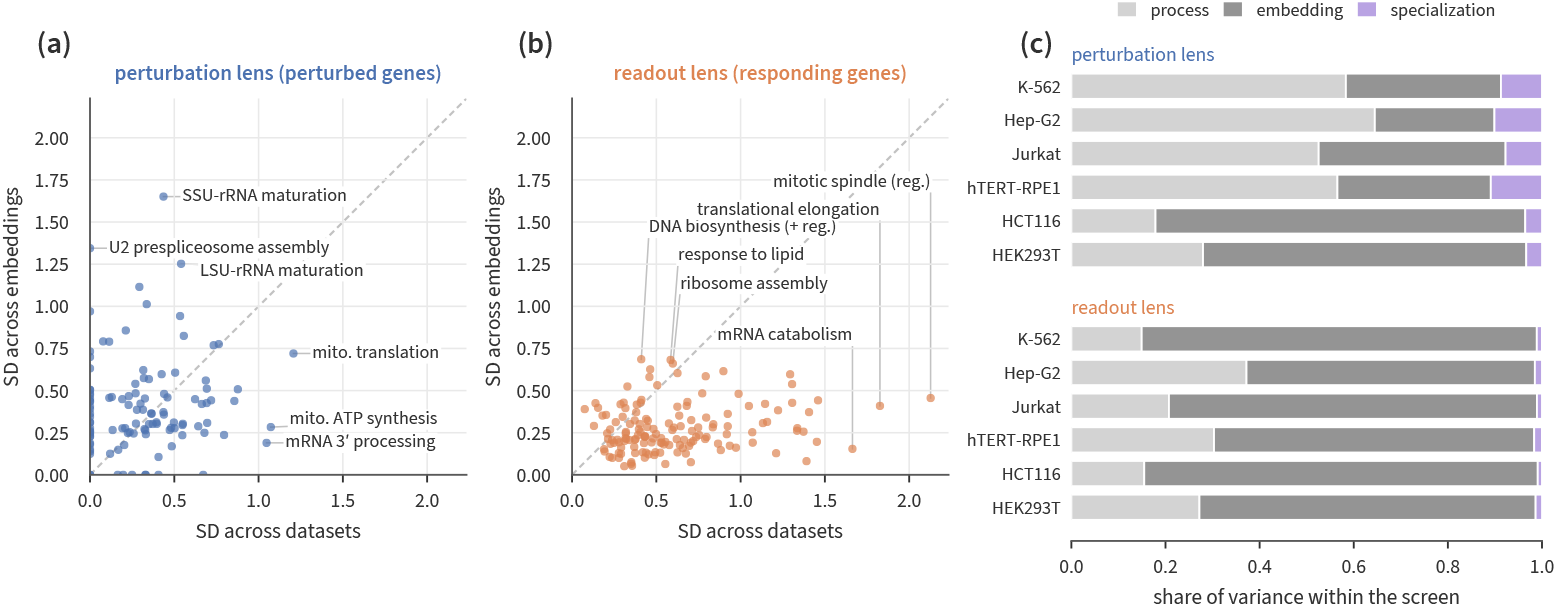
The embedding sets which perturbations are predictable, and the cell line sets which responses are. Predictability of GO biological processes by gene embeddings with lasso in the four Essential and two Xaira screens, scored as R^2^ vs mean (Supplementary Section B.1). The perturbation lens pools R^2^ over a process’s perturbed genes. The readout lens pools it over the process’s genes as responses, on one fixed set of perturbations per screen. (a, b) Each point is a process. The *x* axis is its SD across screens and the *y* axis its SD across embeddings. The dashed line is *y* = *x*. (c) Shares of the within-screen variance of R^2^ due to the process, the embedding, and their interaction (specialization). All quantities are corrected for sampling noise, and corrected SDs below zero are set to zero.

## B Supplemental Methods

### B.1 Metrics

This section defines every metric, gene subset, aggregation, and uncertainty estimate used in the figures. Notation follows Section 4.1.1: Δ_*k*_ ∈ ℝ^*G*^ is the observed log fold-change (LFC) of perturbation *P*_*k*_ over the *G* readout genes (Section 4.1.2) and 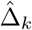 is a model’s out-of-fold prediction. Perturbations are split into five folds. Let K_*j*_ denote the test perturbations of fold *j* and *n*_*j*_ = |K_*j*_|. We write

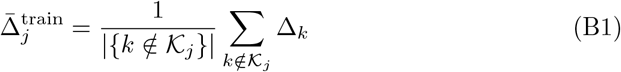

for the mean LFC of the fold’s training perturbations. This is identical to the prediction of the Train Mean baseline.

#### B.1.1 LFC task metrics

##### Traditional regression metrics

For each test perturbation *P*_*k*_ we compute

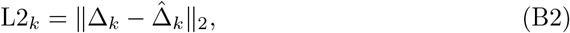

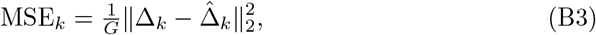

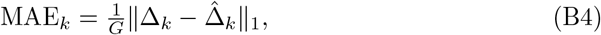

and the Pearson and Spearman correlations between Δ_*k*_ and 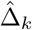 across genes. Pearson and Spearman are undefined when either vector is constant, so No Change 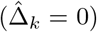 has no correlation score. L2 is the default metric (Section 4).

##### Normalized inverse rank (NIR)

The normalized inverse rank (NIR) asks whether a prediction is closer to its own perturbation’s observed LFC than to the observed LFCs of the other perturbations in the same test fold. For *k* ∈ *K*_*j*_, let

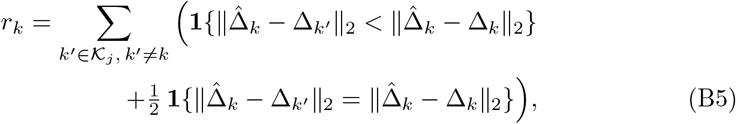

the number of other test perturbations whose observed LFC is closer to the prediction (ties count one half). Then

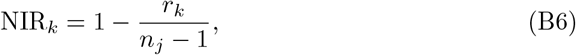

where NIR_*k*_ ∈ [0, 1] and NIR_*k*_ = 1 when the prediction is closest to its own perturbation (higher is better). Because *r*_*k*_ depends on which perturbations share the fold, NIR values are compared between methods on the same folds, and between fold designs (Figure A12) only as differences from the Train Mean (ΔNIR, below).

##### Metrics relative to Train Mean

Figures that report improvement over the Train Mean use paired, per-perturbation differences signed so that positive is better:

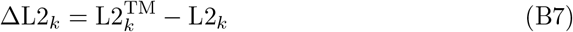

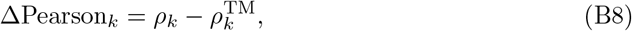

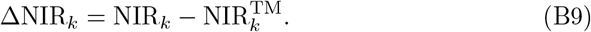

The superscript TM denotes the Train Mean’s score on the same perturbation. The same definitions apply on a gene subset (below), with both scores computed on that subset.

Computing *R*^2^ relative to Train Mean is slightly more involved. This metric is the share of the Train Mean’s squared error that a model removes, pooled over a set *P* of perturbations before division:

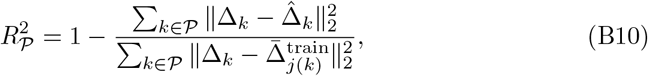

where *j*(*k*) is the fold that holds out *P*_*k*_. 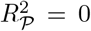 for the Train Mean and 1 for a perfect prediction. Per-method values pool within each fold (*P* =*K* _*j*_) and then average the five folds. The SNR-stratified figures pool over the perturbations of an SNR bin or subset, across folds. The per-perturbation value (*P*= {*k}*) is used only to rank perturbed genes for gene set enrichment (Figure 4).

#### B.1.2 DEG classification

For the DEG task, Δ_*k,g*_ ∈ {−1, 0, +1} (downregulated, no change, upregulated). Models predict per-class probabilities, and the predicted class *ĉ*_*k,g*_ is the most probable one. For fold *j*, gene *g* and class *c*, let 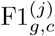 be the F1 score of the predictions 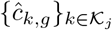 for class *c* against the labels 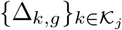, i.e. computed across the fold’s held-out perturbations. Let 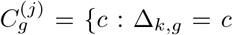 be the classes present in the gene’s labels. The per-gene score is

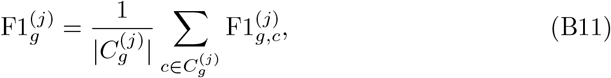

where a present class that is never predicted scores 0 and absent classes are left out (Section B, class imbalance considerations). The fold score averages over the fold’s responsive genes, G_*j*_ = {*g* : Δ_*k,g*_ ≠ 0 for some *k* ∈ K_*j*_}:

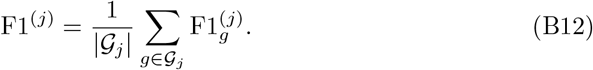

Restricting to responsive genes is intended to keep the score from saturating on genes that are never DE, which may dominate when labels are sparse. The method score is the mean of F1^(*j*)^ over folds.

#### B.1.3 Aggregation

##### Over perturbations and folds

Unless stated otherwise (e.g. for *R*^2^ vs. mean), a method’s score in a fold is the mean of its per-perturbation scores over *K*_*j*_. A fold mean of a correlation is undefined if any perturbation’s correlation is undefined. For the DEG metrics, the model and the Train Mean are averaged over the same set of perturbations, all that have valid scores.

##### Over cell lines

Essential, sci-Plex and the DEG task are reported per cell line. For Tahoe, fold scores are averaged over the 45 cell lines before plotting. The cross-context results average per-cell-line values over the held-out cell lines.

##### Pooled across folds

The SNR-stratified figures (Figures A10, A11, A12) and the fusion bars (Figure 3) pool the out-of-fold per-perturbation scores of all folds and report means over perturbations (or *R*^2^ pooled over them). Each panel is restricted to the perturbations scored by every model it compares; in the SNR-stratified figures, perturbations for which the embedding is missing are dropped.

##### Positive controls

The bootstrap noise oracle (Section 4.7.1) is scored with the same metric as the models: for LFC metrics, the per-perturbation 90th percentile of the bootstrap distribution for errors and the 10th percentile for correlations, then averaged like a model’s scores. Since DEG F1 is a “higher is better” metric, we use the 10th percentile over bootstrap draws of the fold score F1^(*j*)^.

#### B.1.4 Gene subsets

“All gene” metrics use the dataset’s full readout. NIR and *R*^2^ vs mean are reported on all genes only. However, other metrics may be restricted to one of the two gene sets we describe below.

##### DE genes

*D*_*k*_ is the set of readout genes called differentially expressed for *P*_*k*_ by the variance-floored Welch test of the DEG task (Section 4, DEG implementation details), regardless of direction. DE-gene metrics apply the definitions above to 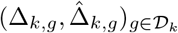. Perturbations without DEG calls (30 or fewer cells) or with |*D* _*k*_| = 0 are excluded from DE-gene metrics. DE-gene L2, MSE and MAE use every perturbation with |D_*k*_| ≥ 1. DE-gene Pearson is reported only for perturbations with |D_*k*_| ≥ 20 to reduce noise. DE-gene Spearman requires only |D_*k*_| ≥ 2 and non-constant vectors.

##### Top-50 SNR genes

S_*k*_ is a fixed-size set of the 50 genes that respond most reproducibly to *P*_*k*_, chosen on held-out cells. Using the split-half construction of Section B.9, each of 10 random splits divides the perturbation’s cells into exact quarters *Q*_0_, …, *Q*_3_, with halves *A* = *Q*_0_ ∪ *Q*_1_ and *B* = *Q*_2_ ∪ *Q*_3_. In each split and each direction (select on *A* or on *B*), genes are scored by 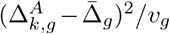, where 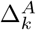 is the LFC estimated from the selection half against all control cells, 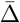 is the mean full-data LFC over perturbations and *v*_*g*_ is the control-cell variance of gene *g* (floored at 10^*−*4^); the perturbed gene itself is excluded, and the top 50 are kept. Selection runs within the 5,539 genes measured in every genetic screen. *S*_*k*_ contains the 50 genes selected most often across the 20 selections, with ties broken by mean selection score. Perturbations for which any gene of *S*_*k*_ is missing from the benchmark readout have no top-50 SNR score.

#### B.1.5 Method rankings

Within each cell line × fold split (e.g. 4 cell lines × 5 folds = 20 splits for Essential), all methods in the figure, including the controls and oracles, are ranked by their fold score: 33 methods for LFC regression (by L2) and 34 for DEG classification (by F1). Rank 1 is best (lowest L2, highest F1), and tied methods receive their average rank. Rows are ordered by mean rank over the 20 splits.

#### B.1.6 Bootstrap confidence intervals

Error bars in the SNR-stratified and fusion figures are 95% percentile intervals from 2,000 bootstrap resamples, with replacement, of the perturbations in the bin, subset or panel. Within a bin or panel, the same resamples are used for every model, so the intervals reflect the sampling of perturbations, not refitting. Paired differences between two methods are computed on the same resamples.

#### B.1.7 SNR stratification

Perturbations are stratified by the split-half SNR of Section B.9 (cell split, each dataset’s own readout). Bins are left-closed, [*a, b*), with edges 0, 0.1, 0.2, 0.3, 0.4, 0.6, 0.8, 1.6, ∞. A bin is shown only if it holds at least 20 perturbations. A DE-gene statistic in a bin is shown only if at least 20 of its perturbations have a value. The “high SNR” subsets of Figure A13 and Figure A12 filter on SNR *>* 0.5. Because SNR sums over the readout genes, it is not comparable across datasets with different readout sets (Section B.9).

#### B.1.8 Metric ranking agreement

To ask whether metrics rank methods consistently (Figure A13), we use 17 metrics, each oriented so that higher is better (errors negated):

- L2, MSE, MAE, *R*^2^ vs mean, NIR, Pearson and Spearman on all genes;
- L2, MSE, MAE, Pearson and Spearman on DE genes; and
- L2, MSE, MAE, Pearson and Spearman on top-50 SNR genes.

For each cell line and regression head (kNN, lasso), each method’s score under each metric is its five-fold mean. We compute the Spearman correlation across methods for every pair of metrics. The matrices are averaged over cell lines and heads (Essential: 4 lines × 2 heads; iPSC: 1 line × 2 heads).

#### B.1.9 Cross-context metrics

The cross-context comparison with STATE (Figure A8) uses the evaluation of Adduri et al. [49], computed with its cell-eval package.

#### B.1.10 Process-level predictability

The perturbation filter, the pooling of *R*^2^ in each lens, the ranking of perturbed genes for enrichment, set selection, the variance decomposition, and its bootstrap correction are described in Section B.8.

### B.2 Embedding Evaluation

#### Embedding preprocessing

In all cases embeddings are standardized and reduced to 100 principal components to control for embedding dimensionality. (Norman is an exception - we use a PCA dimension of 50 due to the smaller number of perturbations in the Norman dataset.)

#### Cross-validation

All four datasets (Essential, Norman, Sciplex, Tahoe) use a nested cross-validation procedure. Within each fold, we perform an inner 5-fold cross-validation with grid search to optimize hyperparameters for the chosen estimator (e.g. kNN, Lasso, Logistic Regression).

#### Splits

In all cases, our splits are defined at the perturbation level. That is, perturbations in the test set have never been seen in any context. In all cases, we generate 5 folds. This means that 20% of perturbations are held out in any given fold. For Essential, Sciplex, Norman, iPSC, and Xaira splits are IID. For Tahoe, we cluster perturbations in terms of their average effect and build stratified splits.

#### Missing embeddings

When an embedding is unavailable from a given source, we simply use the “Train Mean” baseline as our prediction for regression, and “Most Frequent” for classification.

#### B.2.1 Details for LFC Regression

In the LFC regression context, the estimator predicts Δ_*k*_ (Section 4.1.2) and is scored by L2 averaged over perturbations (Section B.1).

##### kNN implementation details

We used the KNeighborsRegressor class from scikit-learn. For Essential, Tahoe, and Sciplex we tune the number of neighbors over the set {20, 40, 60, 80, 100}. For Norman we tune over the set {5, 10, 15, 20}due to its smaller size.

##### Lasso implementation details

We used the Lasso class from scikit-learn. For all datasets (Essential, Tahoe, Sciplex, Norman), we tune the regularization parameter over the set {1e-3, 1e-2, 1e-1, 1}.

#### B.2.2 Details for DEG Classification

In the DEG classification context, the estimator is trained to predict whether each gene is up-regulated, down-regulated, or not significantly changed.

##### Logistic regression implementation details

We implemented Logistic Regression in PyTorch to leverage GPU acceleration. We trained with a weighted cross entropy loss, averaged across gene tasks. Let *ŷ*_*k,g,c*_ be the model prediction for perturbation *k*, gene *g*, and class *c*. We assume that a softmax has been applied over the class dimension, so that *ŷ*_*k,g,c*_ *>* 0 and ∑_*c*_ *ŷ*_*k,g,c*_ = 1. Then the loss is

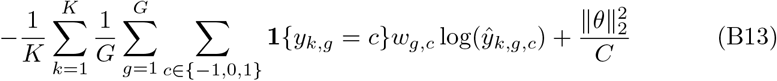

where *θ* is the logistic regression weight matrix and *C >* 0 is a hyperparameter controlling the regularization weight.

The class weighting corresponds to the “balanced” class weighting from scikit-learn. The weight for class *c* and gene *g* is

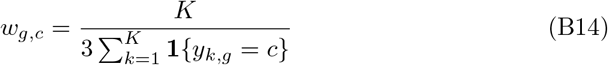

We use the L-BFGS optimizer and tune *C* over the set {1, 1e3, 1e5, 1e7}. This is done independently for each fold and cell line. We handle the Tahoe (which has a large number of cell lines) differently, tuning on a single cell line and using those hyperparameters for all cell lines.

##### Class imbalance considerations

Class imbalance is a challenge in this setting. There are many genes for which a class is never observed in the test set, meaning F1 is undefined (if the class is never predicted) or zero (when the class is predicted at least once).

One could handle this by replacing all NaN values with zeros. However, this shifts all scores toward zero. Even a perfect predictor would be unable to achieve an F1 score of 1.

Another option is to simply ignore NaN scores, excluding them from the macro average. However, this creates an unfair situation: a model that occasionally predicts an absent class would receive a precision of 0 for that class, while a naive model predicting only the majority class would face no penalty. To avoid this inconsistency, we want the set of classes included in metric computation to be independent of model predictions.

We take an intermediate approach by distinguishing between two types of problematic scores: (1) those that can only be NaN or 0 (when a class is absent from the ground truth), and (2) those that can take on a full range of values including NaN, 0, 1, and values in between (when a class exists in the ground truth but may not be predicted). For type (1), we exclude the metric entirely from averaging. For type (2), we convert NaNs to 0, thereby penalizing models that fail to predict classes that do exist in the ground truth.

#### B.3 Advanced Perturbation Prediction Method Details

##### Cross-validation

We perform 5-fold cross-validation for all advanced methods, just as we do for embedding-based methods.

##### Hyperparameter tuning

Due to their high computational cost, it was not feasible to run a large number of Optuna trials for Latent Diffusion, Flow Matching, and Schrödinger Bridge. Instead, we performed a modest amount of manual hyperparameter tuning for each method to arrive at fixed hyperparameters, which we describe below. For GEARS, we performed a grid search over two learning rates (1e-3, 1e-4) and two batch sizes (32, 64) for each fold (holding out 15% of each training fold for validation).

#### B.3.1 Latent Diffusion

For our single cell diffusion model, we encode the samples into a PCA space of dimension 50 by using the PCA basis induced by the training set. Then in this latent space, we train a diffusion transformer (DiT) to learn to predict the noise in the loss function described above. We use adaLN-zero blocks for conditioning, which has been shown in practice to be the best conditioning mechanism for DiTs [82]. We condition our diffusion model on the WaveGC embeddings and train with a constant learning rate of 1e-4 and batch size of 128. We train the models with 5 fold cross validation. For each training fold, we split the perturbations into 85% - 15% and use the 15% validation for early stopping of 20 epochs. The diffusion model learns to transform Gaussian random vectors into perturbed cells conditioned by the WaveGC embeddings. Unlike Flow Matching and Schrödinger Bridge, the Latent Diffusion approach does not use control cells.

Due to training instability there were some cell lines (Hep-G2, Jurkat) that needed to be re-initialized with a different random seed.

#### B.3.2 Flow Matching

For our single cell Flow Matching model, we encode the samples into a PCA space of 128 by taking the PCA loadings of the train set. Then in the latent space, we train the diffusion transformer (DiT) using the Flow Matching objective. Conditioning, learning rate, batch size, cross-validation, and early stopping are as for Latent Diffusion. The Flow Matching model learns to transform control samples into perturbed samples conditioned by the WaveGC embeddings.

#### B.3.3 Schrödinger Bridge

We use the Schrödinger Bridge for single cells in s similar manner to the Flow Matching models. We encode the samples into a PCA space of 50 by taking the PCA loadings of the train set. Then in the latent space, we train the diffusion transformer (DiT) using the Flow Matching objective. Conditioning, learning rate, batch size, cross-validation, and early stopping are as for Latent Diffusion. The Schrödinger Bridge model learns to transform control samples into perturbed samples conditioned by the WaveGC embeddings.

Due to training instability, Fold 2 of the K-562 cell line needed to be re-initialized with a different random seed.

### B.4 Fine-Tuning

#### B.4.1 AIDO.Cell

##### Formulation (Indexing)

Here we describe our approach to fine-tuning an AIDO.Cell model on the LFC regression task. Recall that we aim to predict Δ_*k*_ ∈ ℝ^*G*^, which is defined to be the vector of observed treatment effects for perturbation *P*_*k*_. Let 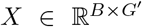 denote a batch of *B* control cell scRNA-seq profiles measured over 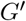 genes. (Note that in general, it is not necessary to have 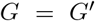.) Let 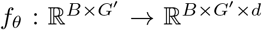 denote the AIDO.Cell encoder (parameterized by *θ*), which consumes a batch of scRNA-seq profiles and produces a *d*-dimensional embedding for each gene and cell. Let’s define a function *m* to compute the mean over the batch dimension and define 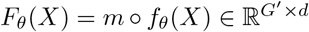. Finally, we introduce a prediction head (e.g. a linear layer or an MLP) *g*_*ϕ*_ : ℝ^*d*^ → ℝ^*G*^. We then define our perturbation response predictor as

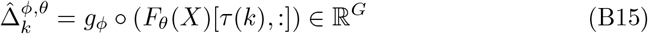

where *τ* (*k*) indexes the gene corresponding to perturbation *P*_*k*_. To fine-tune, we use stochastic gradient descent to optimize

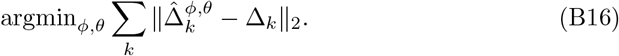

Note that training was performed using data from only a single cell line, so the target output remains fixed regardless of the batch. A natural extension for future work would be to fine-tune across multiple cell lines so the model learns to differentiate perturbation effects in a cell type specific manner.

##### Formulation (In-Silico KO)

Let *y* 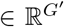 denote the average expression over all control cells. Suppose we want to make a prediction for perturbation *P*_*k*_. Let *τ* (*k*) give the gene index corresponding to perturbation *P*_*k*_. Let 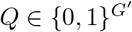 be a masking vector such that *Q*[*i*] = 0 if *i* = *τ* (*k*) and *Q*[*i*] = 1 otherwise. Let 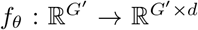 denote the AIDO.Cell encoder (parameterized by *θ*). Let *g*_*ϕ*_ : ℝ^*d*^ →ℝ be a prediction head (parameterized by *ϕ*). Then we compute

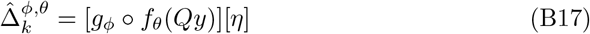

where *η* ∈ ℝ^*G*^ is a set of indices that pick out the genes we want to predict. We then optimize

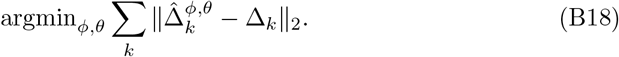

##### Implementation details

For indexing mode, we implement *g*_*ϕ*_ as an MLP with two hidden layers of sizes 1024 and 2048. For in-silico KO, we implement *g*_*ϕ*_ as an MLP with two hidden layers of sizes 64 and 32. We trained using LoRA for 20 epochs for indexing mode and 250 epochs for in-silico KO. Both models seem to have trained to convergence. We use a batch size of 8. In each batch, we randomly select 8 perturbations to update. The only hyperparameter optimization we performed was tuning the learning rate, which we tuned over {1e-2, 1e-3, 1e-4, 1e-5} using the validation set. We explored both the Adam and Muon optimizers but did not observe a significant difference in performance. Within each of the 5 folds, we took 20% of the training data for validation.

#### B.4.2 STRING GNN

##### Formulation

We aim to predict observed treatment effects Δ_*k*_ ∈ ℝ^*G*^ for a perturbation *P*_*k*_. Let *W* denote a graph over *G*^*′*^ genes, with some node embeddings of dimension *d*^*′*^. Let 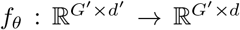 denote our GNN, parameterized by *θ*. Let *g*_*ϕ*_ : ℝ^*d*^ → ℝ^*G*^ be a prediction head, mapping a gene embedding to an estimate 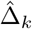 of Δ_*k*_. Define *F*_*θ*_(*k*|*W*) = *f*_*θ*_(*W*)[*τ* (*k*), :] where *τ* (*k*) provides the index corresponding to perturbation *P*_*k*_. Then our prediction for perturbation *P*_*k*_ is

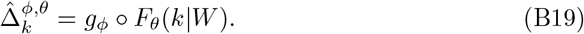

The training objective is

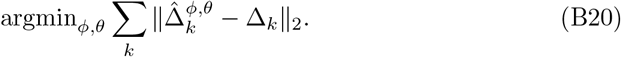

##### Implementation details

We initialize *f*_*θ*_ as the STRING GNN model we used to generate embeddings, which was selected using the validation performance on the link prediction task. We let *g*_*ϕ*_ be a linear projection. We follow the same 5-fold cross-validation splits used throughout the paper for Essential. For each fold, we split off 10% of the training perturbations for a validation set and train on the rest. We tune the learning rate (1e-3, 1e-4, 1e-5) and batch size (16, 32, 64) and select the model with the lowest validation loss. The optimal hyperparameters for each split can be found in Table B1.

### B.5 Full Fusion Model

This section uses the same notation as the description of the simple fusion model in Methods (Section 4).

#### B.5.1 Architecture

For this model, we start by *z*-scoring all of our embeddings using the training set statistics. We will assume this has already been done for *Z*_*k*_. We then compute

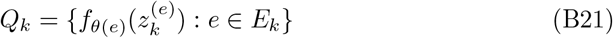

and our input set for perturbation *P*_*k*_ is

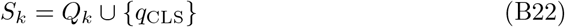

where *q*_CLS_ is a CLS token. Source identity is again left to the bias terms of *f*_*θ*(*e*)_ (Section 4).

**Table B1.** Optimal hyperparameters for each cell line and fold for the fine-tuned STRING GNN model.

| Cell Line | Fold | Batch Size | Learning Rate |
| --- | --- | --- | --- |
| Hep-G2 | 0 | 16 | 1e-5 |
| Hep-G2 | 1 | 16 | 1e-4 |
| Hep-G2 | 2 | 16 | 1e-5 |
| Hep-G2 | 3 | 16 | 1e-5 |
| Hep-G2 | 4 | 16 | 1e-4 |
| hTERT-RPE1 | 0 | 16 | 1e-4 |
| hTERT-RPE1 | 1 | 32 | 1e-4 |
| hTERT-RPE1 | 2 | 32 | 1e-4 |
| hTERT-RPE1 | 3 | 16 | 1e-4 |
| hTERT-RPE1 | 4 | 16 | 1e-4 |
| Jurkat | 0 | 32 | 1e-4 |
| Jurkat | 1 | 16 | 1e-4 |
| Jurkat | 2 | 16 | 1e-5 |
| Jurkat | 3 | 32 | 1e-4 |
| Jurkat | 4 | 16 | 1e-4 |
| K-562 | 0 | 16 | 1e-4 |
| K-562 | 1 | 16 | 1e-4 |
| K-562 | 2 | 16 | 1e-4 |
| K-562 | 3 | 16 | 1e-4 |
| K-562 | 4 | 32 | 1e-5 |

With the transformer *T*_*θ*_ and prediction head *g*_*ϕ*_ of Section 4, we compute *R* = *T*_*θ*_(*S*_*k*_) and predict

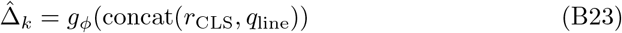

where *r*_CLS_ ∈ *R* is the contextualized output corresponding to *q*_CLS_ and *q*_line_ is the cell line embedding. We then define a training loss *L*_task_(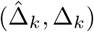, Δ_*k*_), which is either an L2 loss (for LFC regression) or a class-balanced cross-entropy loss (for DEG classification).

Unlike the vanilla transformer used in the simple fusion case, we use gated SDPA (scaled dot product attention) throughout *T*_*θ*_ [83]. This allows the model to dynamically control information flow from each attention operation based on the input context.

The cell line embedding is also more complex in this model. We introduce an autoencoder *h*_*γ*_ : ℝ^*G*^ → ℝ^*G*^ which is trained to reconstruct the average expression of each cell line. We define *q*_line_ to be the latent embedding of *h*_*γ*_. If *y* ∈ ℝ^*G*^ is the mean expression for the cell line of interest, then we train using a standard L2 loss:

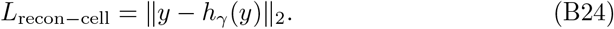

This model also makes use of extra regularizers to prevent overfitting. We use global dropout for all weights and apply dropout to individual input embedding sources to encourage robustness to missing data. Finally, we add a source-specific linear head *W* ^(*e*)^ : ℝ^*d*^ → ℝ^*d*^ which reconstructs the input embeddings from their contextualized variants:

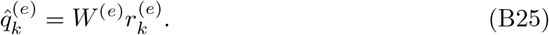

**Table B2.** Optimal hyperparameters for full fusion model tuned on the first fold of the Essential LFC task. These hyperparameters were used for training on folds.

| Parameter | Value |
| --- | --- |
| Learning Rate | 5.28e-4 |
| Batch Size | 32 |
| # Encoder Layers | 8 |
| # Attention Heads | 10 |
| Global Dropout Rate | 0.084 |
| Embedding Source Dropout Rate | 0.004 |
| Prediction Head Depth | 2 |
| Prediction Head Width | 400 |
| $w_{\text{recon-cell}}$ | 9.561 |
| $w_{\text{recon-emb}}$ | 9.412 |

This is done by minimizing an L2 loss:

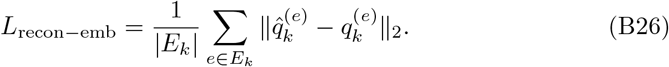

We train the models *T*_*θ*_, *g*_*ϕ*_, {*W* ^(*e*)^}, and *h*_*γ*_ jointly using the combined loss:

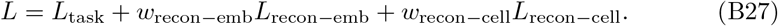

#### B.5.2 Implementation Details

We implement *g*_*ϕ*_ as an MLP. We implement *h*_*γ*_ as an MLP encoder (hidden dimensions 1024, 256) and a decoder MLP (hidden dimensions 256, 1024). We tuned the following hyperparameters: depth (0, 1, 2, 3) and width (200, 400, 800) of *g*_*ϕ*_, number of attention heads (2, 4, 10), number of attention layers (1, 2, 4, 8, 10), batch size (32, 64, 128), learning rate (range from 1e-5 to 9e-3, logarithmic search), global dropout (range from 1e-5 to 0.3, logarithmic search), embedding source dropout (range from 0 to 0.95), *w*_recon*−*cell_ (range from 0.1 to 10), and *w*_recon*−*emb_ (range from 0.1 to 10). We used Optuna with 100 trials, selecting the model with the lowest validation loss on the first fold. Those optimal parameters were then used for all folds. We set *d*_model_ = 100 for *T*_*θ*_. Final hyperparameters can be found in Table B2.

### B.6 Cross-Context Evaluation and STATE Comparison

#### B.6.1 Data

We followed as closely as possible the cross-context formulation from STATE [49] for the Essential and Tahoe datasets.

##### Essential

The Essential dataset was prepared following the notebook^1^ released by the authors of STATE, using a leave-one-out scheme (i.e. holding out one cell line, training on the rest).

##### Tahoe

Starting from the raw Tahoe-100M data [57], we normalized each cell to a total count of 4000 and performed the log1p transformation. Only the highly variable genes (as pre-defined in the STATE paper) were retained. Train, validation, and test splits were also taken from the STATE paper.

#### B.6.2 Models

##### Train Mean

This is a naive baseline method from the STATE paper, where it is called “Context Mean”. To make a prediction for perturbation *P*_*k*_ in an unseen cell line *C*, we simply compute the average response to all perturbations in *C*.

##### kNN

This is a version of kNN regression tailored to the cross-context task, not to be confused with the kNN regression model we use in the unseen perturbation tasks throughout the rest of the paper. To make a prediction for perturbation *P*_*k*_ for a cell from an unseen cell line *C*, we sample a control cell *x* from *C* and identify the *k* = 3 training cell lines whose average control expression has the greatest cosine similarity to *x*. We then average the effect of perturbation *P*_*k*_ across those 3 cell lines and add this to *x*. This simple model allows us to predict different perturbation effects depending on the properties of individual control cells.

##### MLP

We introduce a simple MLP model for mapping a set of control cells to a set of perturbed cells. We now describe the training procedure and model structure. We start by sampling a control cell profile *y* ∈ ℝ^*G*^ and a perturbed cell profile *y*^*′*^ ∈ ℝ^*G*^. Let *x* ∈ ℝ^*d*^ be the embedding of the perturbation received by *y*^*′*^. The MLP *f*_*θ*_ is used to learn a shift that takes *y* to *y*^*′*^. In the genetic perturbation case, we optimize

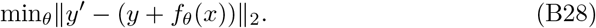

In the chemical perturbation case, we also have to handle dose information. For each dose level *D* in the dataset, we pre-compute a random vector to represent that dose. If *w* ∈ ℝ^*d*^ is the dose embedding for *y*^*′*^, then we optimize

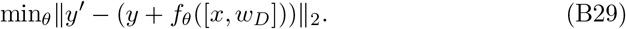

In all cases the MLP *f*_*θ*_ has 3 fully connected layers with ReLU activations. Training was performed using the Adam optimizer with an MSE loss and a batch size of 64. We use a learning rate of 1e-4 for Tahoe and 1e-3 for Essential. No systematic hyperparameter tuning was performed.

By default, the perturbation encoding x is an embedding: WaveGC for the genetic perturbation case (denoted “MLP (WaveGC)” for Essential) and ChatGPT for the small molecule perturbation case (denoted “MLP (ChatGPT)” for Tahoe). As an ablation, we also train a model where *x* is a simple perturbation identity encoding. To do this, we pre-compute a random embedding for each perturbation (similar to our dose encoding scheme) and use those embeddings as *x*. This ablation model is denoted “MLP (One-Hot)”.

##### STATE

We trained STATE [49] (Section 4) from scratch using the official hyperparameter configurations and model code and kept the checkpoint with lowest validation loss. Training from scratch was necessary for two reasons. First, a check-point was not released for the Essential dataset. Second, the official inference tutorial^2^ did not use the same splits as the STATE paper. We judged that retraining was the best way to avoid train/test contamination. We use highly variable genes (HVGs) as input, as STATE does in most of their experiments.

For training we use commit

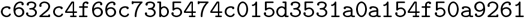

from the official code. For inference, we use commit

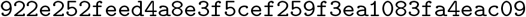

which fixes a critical bug that was causing STATE to produce near-random predictions.

#### B.6.3 Evaluation

We follow the evaluation procedures in STATE, which differ slightly for Tahoe and Essential. For Tahoe the task is zero-shot, so the test set consists of 5 cell lines that are never seen during training. The other 45 cell lines are used for training. The Essential dataset is a few-shot task, where the test set consists of perturbations for one cell line in which a small number of perturbations have been observed. The training and validation is done with all perturbations from three cell lines and a few perturbations from the fourth cell line. Metrics and their aggregation over held-out cell lines are given in Section B.1 (Cross-context metrics).

### B.7 Re-evaluation in the setting of Ahlmann-Eltze et al

For Supplementary Figure A9, we used the single-perturbation benchmark of Ahlmann-Eltze et al. [32] without changes. Its data are the Replogle K562 and RPE1 essential CRISPRi screens [39] as processed by GEARS, with their two train and test splits. A perturbation’s effect is its mean expression minus the control mean. For each embedding, we fit kNN on the training perturbations as in the rest of the benchmark (Supplementary Section B, Embedding Evaluation). The metric is their Pearson delta, the Pearson correlation between predicted and observed effects over the 1,000 genes most expressed in control cells, averaged over test perturbations. Published methods use their reported per-perturbation values. All methods are scored on the test perturbations that every published method scores.

### B.8 Gene set analyses

This section describes Figures A15 and 4. Notation and metric definitions follow Supplementary Section B.1.

#### Data

We used the four Essential screens and the Xaira HCT116 and HEK293T screens, with the benchmark’s out-of-fold lasso predictions. We kept perturbations with replicate SNR above 0.5. The iPSC screen was excluded because only 381 of its perturbations pass this filter. We used the 20 embeddings that beat the Train Mean in all four Essential lines, minus the control PCA and scGPT, which covers under 90% of perturbations. Figure 4 shows one embedding per family.

#### Gene sets

Gene sets come from GO Biological Process 2025 and were chosen without reference to any prediction. A set was a candidate if it held 15 to 500 perturbed genes in at least three screens. We removed redundant sets greedily, from the smallest up. A candidate was dropped if a direct GO edge joined it to a kept set, or if more than half of the smaller set’s perturbed genes lay in the other. This left 144 sets.

#### Two lenses

In the perturbation lens, a set’s score is 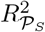, where *P*_*S*_ holds the perturbations of the set’s genes that all 18 embeddings cover. A set is scored if |*P*_*S*_ | ≥ 15. In the readout lens, a set’s score is *R*^2^ over a fixed set of perturbations, restricted to the set’s readout genes. These perturbations are those covered by all 18 embeddings and by scGPT. They are 45% to 92% of the perturbation lens’s perturbations and are biased toward genes expressed in control cells.

#### Spread across screens and embeddings

Scores were centered within each screen and embedding, then scaled within each screen. For each set scored in at least three screens, Figure A15(a, b) plots the standard deviation across screens of its mean over embeddings (x) against the standard deviation across embeddings of its mean over screens (y). These are the main effects of a two-way analysis of variance.

#### Variance shares within a screen

For each screen and lens, we fit an additive model to the table of scores, with sets *s* as rows and embeddings *e* as columns. This gives row effects *a*_*s*_, column effects *b*_*e*_, and residuals *r*_*s,e*_. The process and embedding terms are the sums of squares of *a*_*s*_ and *b*_*e*_. Embeddings that follow the common pattern more or less steeply add a scale term, 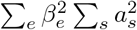, where *β*_*e*_ is the slope of *r*_*s,e*_ on *a*_*s*_ (Finlay–Wilkinson regression). Figure A15(c) counts it with the embedding term. Specialization is the remaining residual sum of squares.

#### Correction for sampling

All statistics in Figure A15 were bias-corrected with 200 bootstrap resamples of perturbations, recomputing every step. The corrected value is twice the observed value minus the bootstrap mean, floored at zero for variances. In the perturbation lens, sets were resampled independently, which ignores covariance between sets that share perturbed genes.

#### Enrichment analysis

For each screen and embedding, we ranked the covered perturbed genes by 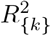 and ran preranked GSEA (gseapy, 1,000 permutations) on the 144 sets. A positive normalized enrichment score (NES) means the set’s perturbed genes are predicted better than the embedding’s others. Figure 4 shows every set with FDR *<* 0.05, in either direction, for at least one shown embedding and screen. Rows are grouped by GO ancestry.

### B.9 Per-Perturbation Signal-to-Noise Ratio

Some perturbations may have a strong effect, while others may be indistinguishable from noise. We compute a per-perturbation split-half signal-to-noise ratio (SNR) to measure this property. This section describes the procedure for calculating this quantity.

Suppose we begin with a population of control cells and a population of cells that received some perturbation *k*. Let Δ_*k*_ be the estimated average treatment effect for perturbation *k*, and let 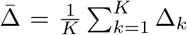 be the average response over all perturbations. We model 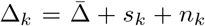 where *s*_*k*_ is the perturbation-specific signal and *n*_*k*_ is the mean-zero perturbation specific noise. Then we can compute

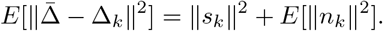

To estimate ∥*s*_*k*_∥ ^2^, we will randomly split the perturbed cells into two equal halves. We do this *R* = 50 times, and index the repetitions by 1 ≤*r* ≤*R*. For each half we compute the treatment effect using the same estimator as Δ_*k*_, which we call 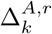 and 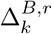. Note that all control cells are used to define the control response, i.e. control cells are not split. If both halves estimate the same signal *s*_*k*_ with independent zero-mean additive noise, then

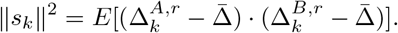

which we estimate as

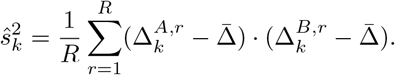

We know 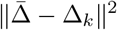 already, so we can plug in

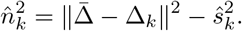

Finally, we can compute

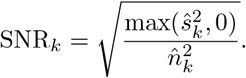

Recall that we compute the ground truth average treatment effect differently depending on the dataset, as described in Section 4.1.2. For a given dataset, we use the same treatment effect calculations to define the ground truth and to compute SNR values.

It is also important to keep in mind that we are treating as “signal” whatever is reproducible in both halves of a random split. This signal likely combines biological and technical effects. Thus, the SNR for the true biological signal is likely to be lower than our estimates.

**Table B3.** Hyperparameters for noise oracle calculations. Note that the batches for Essential are too small for per-batch differential expression, so we ignore batches for the DEG Classification setting.

| Dataset | Setting | $M_{\text{outer}}$ | $M_{\text{inner}}$ |
| --- | --- | --- | --- |
| Essential | LFC Regression | 10 | 10 |
| Essential | DEG Classification | - | 20 |
| Tahoe | LFC Regression | 10 | 20 |
| Tahoe | DEG Classification | 3 | 3 |
| Sciplex | LFC Regression | 10 | 20 |
| Sciplex | DEG Classification | 3 | 3 |
| Norman | LFC Regression | 10 | 10 |

Another caveat is that the SNR depends on the set of genes under consideration, since the inner products and norms are computed over genes. We use each dataset’s own set of readout genes. For this reason, SNR values are not comparable across datasets.

### B.10 Oracles

#### B.10.1 Noise Oracle (Bootstrap)

We show the hyperparameters for noise oracle calculations in Table B3.

### B.10.2 Half-Split Noise Oracles

Here we describe two half-split noise oracles: cell split and GEM split. Both follow the split-half construction of Section B.9: *R* = 50 random halves per perturbation, estimates 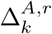 and 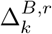 from each half against the unsplit control population, and the agreement between the halves as the estimate of the error of a predictor that knows the true effect. They differ in how the halves are drawn. The cell split assigns cells at random, and the GEM split assigns GEM groups at random.

Following a similar argument as in Section B.9, we can estimate the squared norm of the true treatment effect as

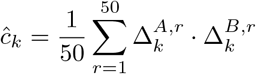

from which we define the oracle value:

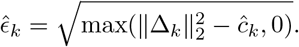

### B.11 Perturbation Source Similarity

Figure 2 is a clustered heatmap plot of similarity scores between different embedding sources. The similarity between a pair of embedding sources is defined to be the mean squared canonical correlation, which is computed via Canonical Correlation Analysis (CCA). We restrict all embedding collections to only include those genes that are present in every embedding, leaving 5348 genes. We then perform PCA to reduce dimensionality to 10 before performing CCA. PCA is performed independently for each embedding source. We adapt the code from Zhong et al. [84] to perform our analysis.

### B.12 Small Molecule Embeddings

#### B.12.1 Strategies for Embedding Small Molecules

Figure B16 visualizes different small molecule embedding strategies.

#### B.12.2 LLM Prompts for Target Identification

Below we include a sample prompt for our LLM-based target identification approach:

~~~
You are a pharmacology expert. For the compound ‘{compound}’,
provide exactly 3 most well-established gene targets in JSON format:
{{‘compound’: ‘{compound}’, ‘targets’: [{{‘gene’: ‘GENE NAME’, ‘confidence’:
‘high/medium/low’, ‘mechanism’: ‘brief description’}}]}}
~~~

#### B.12.3 Affinity-Based Embeddings

The fragment-based embeddings are computationally cheaper because co-folding small molecules with short sequences is much faster than with longer protein sequences. The full protein embedding is more expensive but may more realistically reflect interactions that could occur in the cell, recover protein targets, and capture off-target effects.

##### Fragment-based embeddings

The fragment embedding uses a set of 1000 protein fragments. We selected these fragments with the help of the CATH database, a resource that provides information on the evolutionary relationships of protein domains. The database was used to annotate all domains of human proteins in the Protein Data Bank. We identified the 1000 CATH classes that appear most frequently in the human proteome and select the first representative provided by CATH for each class (i.e. a more or less arbitrary representative).

**Fig. B16.**
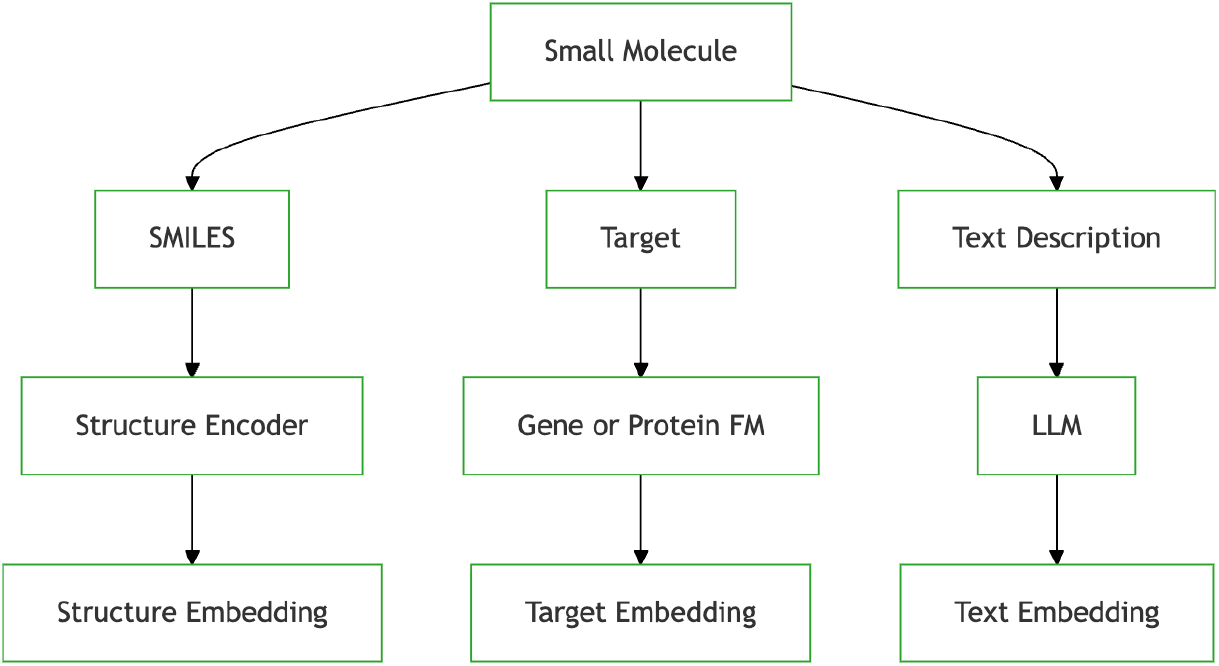
Small molecules can be represented by embedding their structure, their targets (if available), or text describing the molecule (if available).

##### Protein-based embeddings

The protein embedding uses 800 rule-selected targets. First, we obtain disease relevance score from OpenTargets [85] which we use as a proxy for biological importance. We then annotate proteins with their ChEMBL protein classes and with unsupervised cluster labels based on sequence similarity computed with MMseqs2 [86]. We use these categories to ensure diversity in the set of protein targets.

A protein is included as a target if it meets either of the following criteria: (i) the protein is among the 11 top-scoring in its ChEMBL cluster and is the highest-scoring in its sequence similarity cluster; or (ii) the protein is annotated as a target in the Tahoe-100M dataset.

## Footnotes

1 https://colab.research.google.com/drive/1Ih-KtTEsPqDQnjTh6etVvf-gRAA86ZN

2 https://colab.research.google.com/drive/1bq5v7hixnM-tZHwNdgPiuuDo6kuiwLKJ

## Notes

https://github.com/genbio-ai/foundation-models-perturbation

### Competing Interest Statement

All authors are affiliated with GenBio AI.

### Summary of Updates

All figures redrawn; text revised and expanded; additional results and new figures added; supplemental results no longer kept in a separate file.

